# Long-term voluntary exercise reveals limited translation of hippocampal molecular responses into neuroprotection in 5xFAD mice

**DOI:** 10.64898/2026.09.16.752181

**Authors:** Karel Aceituno, Katrina Granger, Jocelyne Leon, Jose A. Godoy-Lugo, Khristina E. Young, Tiarra Joseph, Kailin Liu, Iris Kruijff, Emily Morales, Mia Hakian, Allison Birnbaum, Rik van der Kant, Cristal M. Hill, Constanza J. Cortes

## Abstract

Physical exercise promotes systemic and neural adaptations that support healthy brain aging and may mitigate Alzheimer’s disease (AD) progression. However, the capacity of the AD-afflicted brain to mount and translate exercise-responsive molecular adaptations into neuroprotection remains unclear. Here, we examined the effects of long-term voluntary wheel running (VWR) on molecular, neuropathological, and behavioral outcomes in independently studied male and female 5xFAD mice. VWR elicited expected metabolic and transcriptional remodeling of inguinal white adipose tissue, confirming engagement of exercise-responsive peripheral biology. In contrast, hippocampal transcriptional responses were modest, with few differentially expressed genes and coordinated changes emerging primarily at the pathway level. These responses involved synaptic, neuroimmune, mitochondrial, neurotrophic, and monoaminergic processes and differed qualitatively between the two groups. Several components of the canonical hippocampal exercise response also failed to converge into coordinated cellular adaptations: synaptic protein abundance changed without altering synapse density, while neurotrophic, neurogenic, and vascular responses showed little correspondence across molecular and cellular measures. VWR also produced little change in hippocampal amyloid pathology or behavioral function despite sustained exercise engagement. Together, these findings demonstrate that the 5xFAD brain retains modest molecular responsiveness to prolonged voluntary exercise but may be unable to mount a sufficiently robust or coordinated response to produce broad neuroprotective effects. These findings highlight disease context as an important determinant of the efficacy of exercise-based interventions in neurodegenerative disease.

## INTRODUCTION

Physical activity exerts broad effects on brain health ^1,2^ and remains one of the modifiable factors most consistently associated with reduced risk of age-related cognitive decline and dementia ^3–7^. Endurance exercise (such as running) promotes hippocampal plasticity through multiple mechanisms, including enhanced neurotrophic signaling, re-activation of adult neurogenesis, promotion of synaptic remodeling, vascular adaptations and mitochondrial function, as well as modulation of neuroimmune activity ^1,8,9^. These responses have motivated substantial interest in exercise as a non-pharmacological strategy to preserve cognitive function and potentially modify the progression of Alzheimer’s disease (AD). Consistent with this premise, exercise improves cognitive and neuropathological outcomes across multiple preclinical AD models, including attenuation of amyloid and tau pathology and neuroinflammation, together with enhanced synaptic and neurogenic function ^10–14^.

Large-scale and cell-resolved transcriptomic studies demonstrate extensive molecular remodeling across the healthy brain and peripheral tissues following exercise ^9,15,16^, yet considerably less is known about how established AD pathology alters this adaptive response or whether residual molecular plasticity predicts downstream structural and functional benefit. Indeed, despite the predominantly beneficial effects reported across the preclinical literature, exercise does not uniformly confer neuroprotection in AD models, with interventions producing robust cognitive and neuropathological benefits under some conditions ^10–14^ and limited or absent effects under others ^17–19^. Similar heterogeneity occurs clinically across the disease continuum, from mild cognitive impairment (MCI), a symptomatic stage associated with increased risk of progression to AD dementia, to established AD, with exercise interventions producing variable cognitive and cerebral outcomes^20–22^. Several factors may contribute to these variable responses, including disease stage and severity ^17–19,23^, exercise modality, and the resulting differences in exercise intensity, workload, and physiological and metabolic stress ^24–26^.

These considerations are particularly relevant to the 5xFAD transgenic mouse model, which develops rapidly progressive amyloid pathology accompanied by neuroinflammation, synaptic dysfunction, and behavioral abnormalities ^27,28^. 5xFAD mice have been widely used to evaluate candidate disease-modifying interventions ^28^, including exercise ^11,12,17,19^, yet exercise responsiveness within this model remains strikingly variable. Voluntary wheel running (VWR) has improved cognition, neurotrophic signaling, neurogenesis, and amyloid-related pathology in some studies ^11,12^, whereas others report minimal effects on neuropathology and behavioral outcomes ^17,24^. Moreover, voluntary running elicited markedly attenuated transcriptional remodeling in the 5xFAD brain compared with wild-type mice despite similar running performance ^19^, raising the possibility that the 5xFAD disease environment constrains the molecular plasticity normally elicited by exercise. Together, these findings raise an important question of whether prolonged voluntary exercise can sufficiently engage neuroprotective pathways to modify disease-associated outcomes in the context of rapid progressive amyloid pathology. Moreover, the extent to which the 5xFAD hippocampus retains molecular responsiveness to prolonged VWR, and whether these adaptations translate into cellular and functional benefits, remains unclear.

Here, we investigated the effects of long-term VWR on hippocampal molecular and cellular plasticity, AD-related neuropathology, and behavioral function in independently studied male and female 5xFAD cohorts. We integrated hippocampal transcriptomics with targeted protein and histological analyses to determine whether exercise-responsive transcriptional programs corresponded with downstream cellular adaptations.

Because exercise elicits coordinated adaptations across central and peripheral tissues, we also assessed physiological and transcriptional remodeling of inguinal white adipose tissue (iWAT), a metabolically plastic peripheral tissue that undergoes extensive adaptation in response to exercise ^29,30^. Long-term VWR elicited the expected transcriptional adaptations in iWAT, confirming cellular/molecular/genetic engagement of the exercise intervention, while the hippocampus showed limited global transcriptional remodeling but detectable changes in discrete biological pathways. Thus, these molecular changes occurred without broad modification of amyloid pathology, neurogenesis, or cognitive function. Together, these findings suggest that established 5xFAD pathology may constrain the translation of VWR-responsive molecular signals into coordinated neuroprotective adaptations, underscoring the importance of disease context when evaluating the neuroprotective potential of exercise interventions.

## RESULTS

### Long-term voluntary wheel running elicits physiological adaptations in 5xFAD mice

We ^31,32^ and others ^12,33–35^ have demonstrated that voluntary wheel running (VWR) promotes both peripheral and CNS metabolic and functional adaptations and can modify pathological findings and improve neurocognitive performance in models of neurodegenerative disease ^12–14^. To determine the effects of long-term voluntary exercise in the 5xFAD transgenic line, heterozygous 5xFAD^+^ male and female mice were provided continuous, individual access to either locked (sedentary) or free (running) voluntary running wheels (**Fig. 1A** and **S1A**). Because female 5xFAD mice develop amyloid pathology earlier and more rapidly than males ^27,28^, we initiated VWR at ∼3 months of age in females and ∼6 months of age in males to better align amyloid disease burden across cohorts. Both cohorts underwent 16 weeks of VWR before endpoint analyses between 7-8 and 11-12 months of age, respectively. Consistent with this design, sedentary 5xFAD males and females showed comparable hippocampal Aβ plaque number and astrocyte/microglial morphometrics (as a proxy for neuroinflammatory states) at endpoint, although total plaque volume remained lower in females (**Supplementary Fig. 1B, C**). Given the different ages of the male and female cohorts, throughout this work we evaluated the effects of VWR relative to age-matched sedentary controls within each sex and avoided direct comparisons of exercise responses between sexes.

**Fig. 1.**
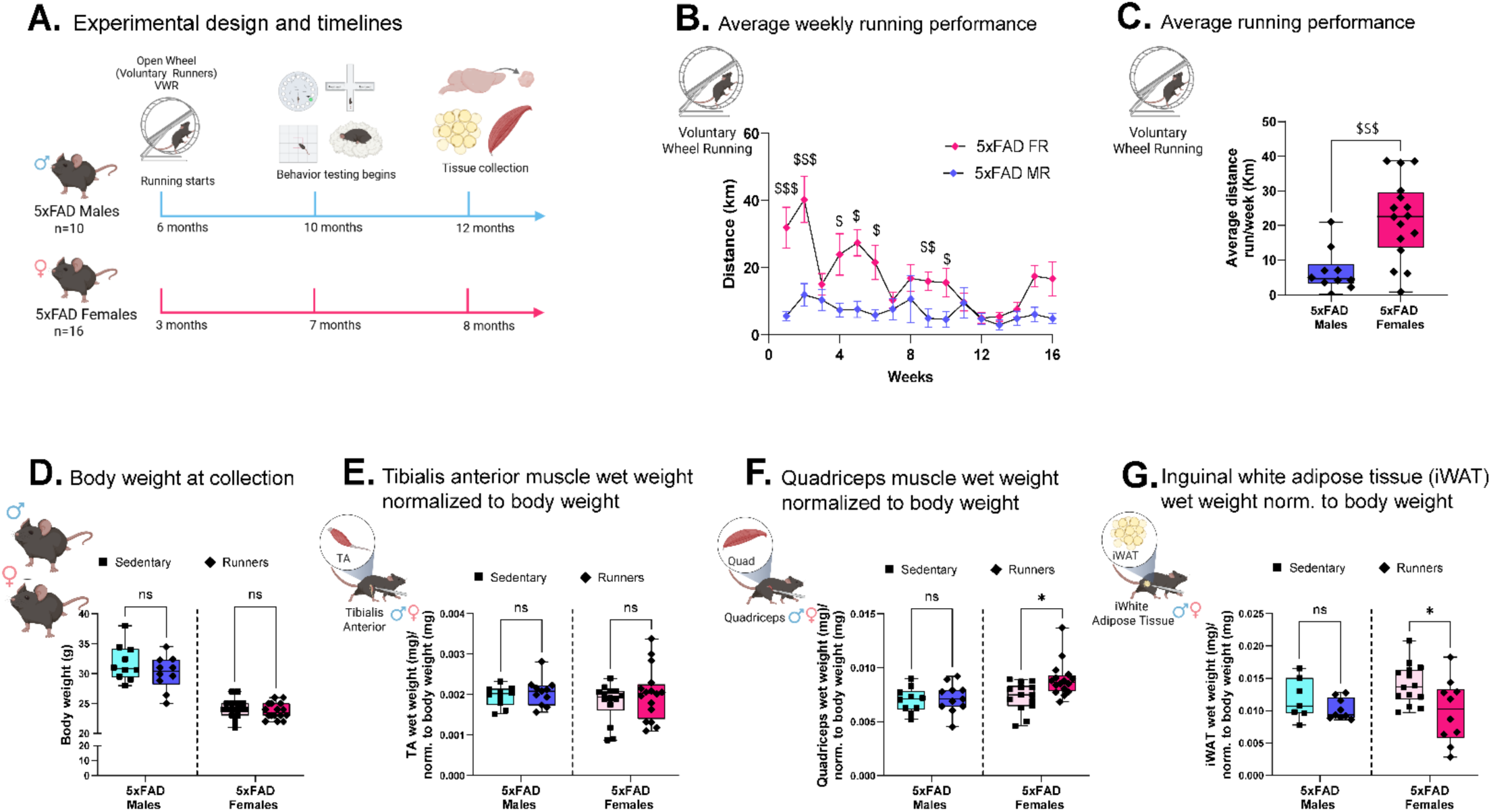
Long-term voluntary wheel running elicits physiological adaptations in 5xFAD mice. **A.** Experimental design. Female 5xFAD mice began voluntary wheel running (VWR) or sedentary locked-wheel exposure at 3 months of age and male 5xFAD mice at 6 months of age; both cohorts underwent 16 weeks of intervention before endpoint analyses. **B.** Weekly running distance throughout the 16-week intervention in male and female VWR cohorts. **C.** Average weekly running distance. **D.** Body weight at endpoint. **E.** Tibialis anterior wet weight normalized to body weight. **F.** Quadriceps wet weight normalized to body weight. **G.** Inguinal white adipose tissue (iWAT) wet weight normalized to body weight. Data are presented as mean ± SEM. Each point represents one animal. n = 10-16 animals/group. Asterisks denote comparisons between sedentary and VWR groups within each cohort (unpaired two-tailed Student’s t-test); dollar signs denote comparisons of running distance between the independently studied male and female VWR cohorts (B. two-way repeated-measures ANOVA, C. unpaired two-tailed Student’s t-test). *,^$^ P < 0.05, ^$$^ P < 0.01, ^$$$^ P < 0.001.

We first confirmed that both male and female 5xFAD mice engaged in sustained voluntary running throughout the entire 16-week intervention (**Fig. 1B**). Female runners averaged approximately twice the weekly running distance of male runners (**Fig. 1C**), consistent with the higher voluntary running distances typically observed in female wild-type mice of similar ages ^29,36^. Despite prolonged VWR, total body weight did not differ between sedentary and runner mice within either the male or female cohort at the end of the experiment (**Fig. 1D**). To determine whether VWR elicited peripheral physiological adaptations, we next assessed skeletal muscle and inguinal white adipose tissue (iWAT) wet weight, two peripheral tissues that undergo substantial remodeling in response to exercise training. VWR did not alter skeletal muscle tibialis anterior mass normalized to body weight in either cohort (**Fig. 1E**). Similarly, quadriceps mass did not differ between sedentary and runner males, whereas female runners exhibited slightly increased quadriceps mass relative to age-matched sedentary females (**Fig. 1F**). VWR also reduced normalized iWAT mass in female runners, while iWAT mass remained unchanged in the male cohort (**Fig. 1G**). Together, these findings demonstrate that long-term VWR is (a) tractable in 5xFAD mice, (b) elicits measurable peripheral physiological adaptations, and (c) produces distinct patterns of peripheral remodeling in the independently studied male and female cohorts.

### VWR induces selective synaptic and mitochondrial adaptations in the male 5xFAD hippocampus

To determine how long-term VWR alters the molecular landscape of the 5xFAD hippocampus, we isolated RNA from the hippocampus of male sedentary and runner 5xFAD mice and performed bulk RNA sequencing (RNA-seq, n=5/group). Principal component analysis (PCA) revealed substantial overlap between sedentary and runner hippocampal transcriptomes, indicating that VWR did not induce broad separation of the two groups at the global transcriptomic level (**Fig. S2A**). Consistent with this pattern, differential expression analysis identified only 4 transcripts significantly downregulated by VWR after multiple-testing correction: *Ttr*, *Slc4a5*, *Igfbp2*, and *Elovl7* (FDR < 0.10 and |log2(fold change)| ≥ 0.58, **Fig. S2B**). Despite the limited differential expression at the individual-gene level, Gene Set Enrichment Analysis (GSEA) identified coordinated pathway-level responses to VWR by detecting consistent shifts among functionally related genes across the ranked transcriptome, even when individual genes did not meet differential-expression thresholds. Across GO domains, GSEA identified 176 Biological Process (BP), 34 Cellular Compartment (CC), and 29 Molecular Function (MF) gene sets that met the significance threshold (FDR < 0.05). The top 10 positively and negatively enriched gene sets from each domain are shown in **Fig. 2A** and **Fig. S2C–D**, with complete GSEA results provided in **Supplementary Table 1**. We focused subsequent analyses on biological programs that showed convergence across GO domains and/or had established relevance to exercise-responsive hippocampal biology.

**Figure 2.**
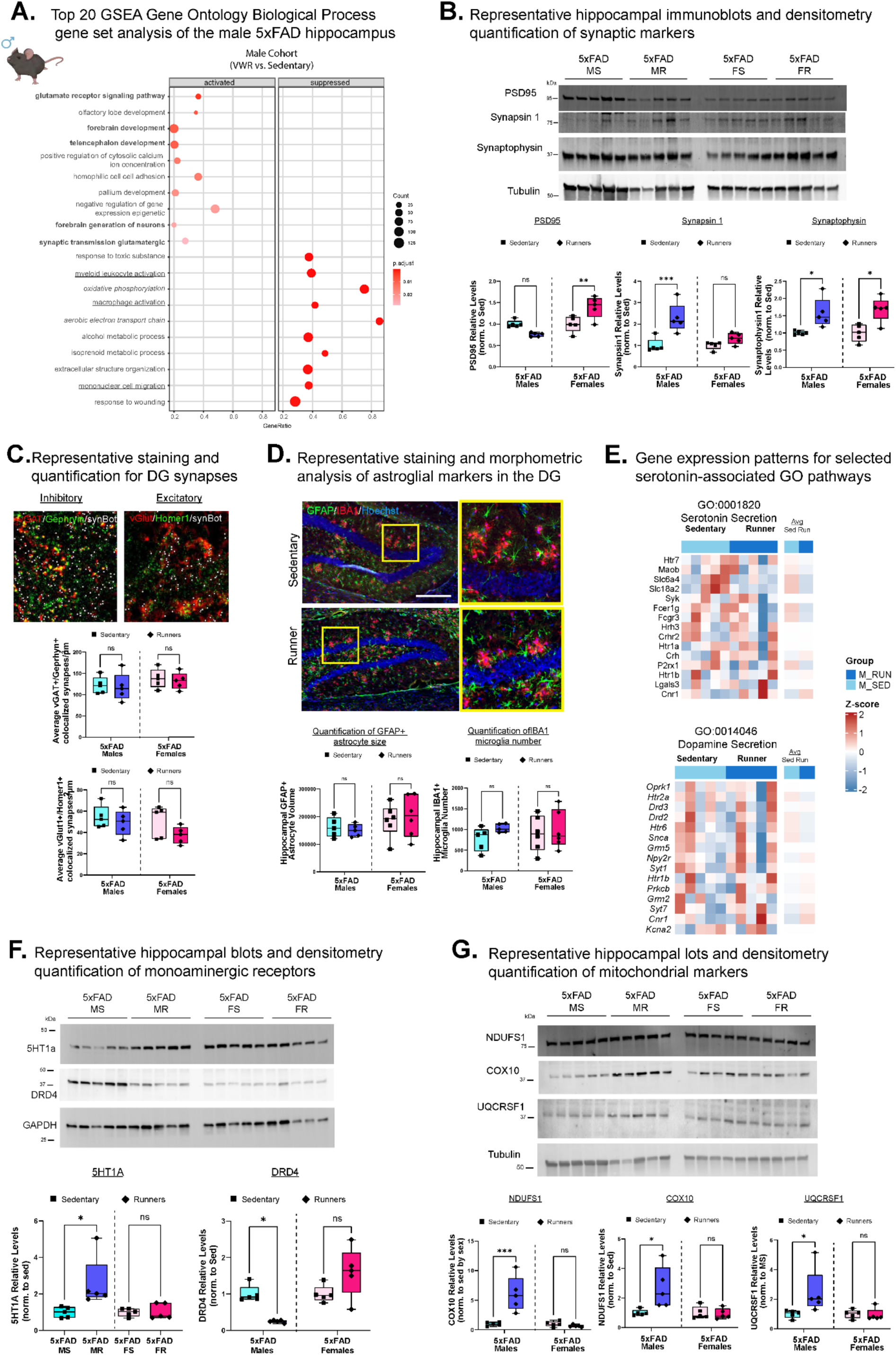
Long-term voluntary wheel running elicits modest synaptic, glial, monoaminergic, and mitochondrial remodeling in the 5xFAD hippocampus. **A.** Gene Set Enrichment Analysis (GSEA) of Gene Ontology Biological Process (GO:BP) terms comparing hippocampi from sedentary and VWR male 5xFAD mice. The 10 most positively and negatively enriched gene sets are shown; complete enrichment results are provided in Supplementary Table 1. **B.** Representative immunoblots and quantification of the synaptic proteins in hippocampal lysates from male and female cohorts. **C.** Representative dentate gyrus immunofluorescence and SynBot quantification of inhibitory and excitatory synapses. **D.** Representative GFAP and IBA1 immunofluorescence and quantification of astrocytic and microglial endpoints in the male dentate gyrus. Scale bars, 100 μm. **E.** Heatmaps showing normalized expression of genes (z-scores) comprising representative serotonin- and dopamine-associated GSEA gene sets in the male hippocampus. Columns represent individual animals followed by group-averaged expression for sedentary and VWR groups. **F.** Representative immunoblots and quantification of hippocampal 5-HT1A and DRD4 protein abundance in male and female cohorts. **G.** Representative immunoblots and quantification of the mitochondrial respiratory chain-associated proteins in male and female hippocampal lysates. Unless otherwise noted, data are presented as mean ± SEM, with each point representing one animal. n = 5-6 animals/group. n.s., not significant; *P < 0.05, **P < 0.01, ***P < 0.001, unpaired two-tailed Student’s t-tests.

Among the significantly enriched GO:BP terms, VWR positively enriched pathways associated with several neuronal and synaptic processes, including *glutamatergic synaptic transmission* and *glutamate receptor signaling* (**Fig. 2A**, in bold). Consistent with these findings, analysis of GO Cellular Component terms identified positive enrichment of *excitatory synapse*, *postsynaptic density membrane*, *ionotropic glutamate receptor complex*, *postsynaptic specialization membrane*, and *GABAergic synapse gene sets* (**Fig. S2C,** in bold). GO Molecular Function analysis similarly identified positive enrichment of *glutamate receptor activity*, *ligand-gated calcium channel activity*, and *calcium ion transmembrane transporter activity* (**Fig. S2D,** in bold). Together, these complementary enrichment analyses identified synaptic and neurotransmission-related processes as prominent components of the VWR response in the male 5xFAD hippocampus. We next used these transcriptomic findings to guide targeted molecular and cellular analyses, examining selected endpoints in both cohorts to determine whether the associated exercise responses extended beyond the cohort in which they were initially identified.

To determine whether this transcriptional remodeling extended to synaptic protein abundance, we quantified the postsynaptic scaffolding protein PSD95 and the presynaptic proteins Synapsin 1 and Synaptophysin in hippocampal lysates (**Fig. 2B**). VWR had no effects PSD95 abundance but increased Synapsin 1 and Synaptophysin in males. The independently studied female cohort also showed increased PSD95 and Synaptophysin, with no change in Synapsin 1 (**Fig. 2B**). We next determined whether these changes in synaptic protein abundance reflected alterations in synapse density. Using SynBot, an open-source image-analysis tool for automated synapse quantification ^37^, we analyzed hippocampal immunofluorescence for colocalized pre- and postsynaptic puncta (**Fig. 2C**). We defined inhibitory synapses by vGAT^+^–Gephyrin^+^ colocalization and excitatory synapses by vGLUT1^+^–Homer1^+^ colocalization. VWR did not alter inhibitory or excitatory synapse density in the dentate gyrus of either cohort examined (**Fig. 2C**). Thus, changes in synaptic protein abundance occurred without detectable remodeling of synapse density in the dentate gyrus of the 5xFAD hippocampus.

In contrast to the positively enriched neuronal and synaptic pathways, VWR suppressed several immune-related GO:BP processes, including *myeloid leukocyte activation*, *mononuclear cell migration*, and *macrophage activation* (**Fig. 2A**, underlined). Because microglia represent the resident myeloid population of the CNS and astrocytes contribute broadly to the neuroinflammatory response, we next examined whether these transcriptional signatures were associated with altered glial responses. At baseline, and despite their different chronological ages, sedentary 5xFAD males and females showed comparable astrocyte and microglial morphometrics (**Fig. S1C**). We therefore performed immunofluorescence staining to assess astrocytic (GFAP^+^) and microglial (IBA1^+^) abundance and morphology, parameters associated with changes in glial activation and inflammatory state (**Fig. 2D**). VWR did not significantly alter the average size or number of GFAP^+^-objects in the male 5xFAD dentate gyrus (**Fig. 2D; Fig. S2E**). Analysis of the distribution of GFAPpositive structures across size ranges similarly revealed comparable profiles between sedentary and runner males (**Fig. S2F**). VWR also did not alter the number of IBA1-positive microglia in the 5xFAD male dentate gyrus (**Fig. 2D**). To further assess microglial morphology, we classified IBA1-positive cells according to morphological states ranging from steady-state and hyper-ramified to amoeboid morphologies (**Fig. S2G**). The relative distribution of these morphological states was similar between sedentary and runner males (**Fig. S2H**), with mild to no changes in morphological state distribution in females. Together, these findings indicate that detectable changes in the measured astrocyte or microglial abundance and morphometric endpoints did not accompany the immune-related transcriptional changes identified by GSEA.

GSEA also identified exercise-responsive pathways related to monoaminergic neurotransmission in the male 5xFAD hippocampus (**Supplementary Table 1**). To examine the gene-level patterns underlying these pathways, we plotted normalized expression of genes comprising representative serotonin- and dopamine-associated gene sets, including group-averaged expression for sedentary and VWR groups (**Fig. 2E**). These heatmaps revealed coordinated but heterogeneous expression changes across genes involved in monoaminergic signaling, consistent with pathway-level remodeling rather than uniform regulation of individual transcripts. Notably, independent phenotype enrichment using the Monarch Initiative similarly identified serotonin- and dopamine-associated phenotypes among exercise-responsive genes (data not shown), further supporting monoaminergic signaling as a component of the 5xFAD male hippocampus transcriptional response to VWR. Given this enrichment of phenotypes associated with altered serotonin and dopamine signaling, we next asked whether VWR altered hippocampal serotonergic and dopaminergic receptor abundance. We quantified 5-HT1A, a serotonin receptor with prominent roles in hippocampal serotonergic signaling, and DRD4, a dopamine receptor that modulates neuronal excitability and synaptic transmission, by immunoblotting of total hippocampal lysates (**Fig. 2F**). VWR increased total 5-HT1A protein levels and decreased DRD4 protein abundance in male 5xFAD mice, with neither receptor altered in the female cohort (**Fig. 2F**). These findings provide evidence of exercise-responsive remodeling of serotonergic and dopaminergic signaling components in the male 5xFAD hippocampus.

Negative enrichment following VWR also extended to mitochondrial pathways, with multiple gene sets related to *oxidative phosphorylation* and *mitochondrial electron transport* emerging from the GSEA (**Fig. 2A**, in italics). This finding was notable given the well-established mitochondrial dysfunction in AD and the importance of mitochondrial remodeling in the response to exercise ^38^. This pattern extended across GO domains, with VWR negatively enriching *mitochondrial respiratory chain* and *cytochrome complexes* in GO:CC (**Fig. S2C,** in italics) and *electron transfer activity* and *oxidoreduction-driven active transmembrane transporter activity* in GO:MF (**Fig. S2D,** in italics). To determine whether these transcriptional signatures extended to the protein level, we quantified proteins associated with mitochondrial respiratory chain complexes I (NDUFS1), III (UQCRFS1), and IV (COX10) in total hippocampal lysates by immunoblotting (**Fig. 2G**). Despite the negative enrichment of mitochondrial gene sets observed in the GSEA, VWR increased NDUFS1, COX10, and UQCRFS1 protein abundance in the hippocampus of male 5xFAD mice relative to their age-matched sedentary controls. VWR did not alter any of the three mitochondrial proteins examined in the hippocampus of the female cohort. This discordance suggests that transcriptional changes may not directly predict mitochondrial protein abundance following VWR.

Together, these findings demonstrate that long-term VWR produced a modest but coordinated pathway-level response in the male 5xFAD hippocampus, characterized by transcriptional remodeling of synaptic, immune-related, mitochondrial, and neurotransmitter-related pathways and discrete changes in associated proteins, but limited changes in cellular inflammatory endpoints. Assessment of these same molecular and cellular endpoints in the independently studied female cohort revealed only partial concordance, with some exercise-responsive features shared across cohorts and others observed only in males.

### VWR alters immune and neuronal transcriptional programs in the female 5xFAD hippocampus

We next applied the same transcriptomic analysis pipeline to the 5xFAD female cohort (n=5/group). As in males, PCA showed substantial overlap between hippocampal transcriptomes from sedentary and runner females, indicating limited global transcriptional separation following VWR (**Fig. S3A**). Consistent with this pattern, no individual transcripts met the predefined differential-expression criteria outlined above (**Fig. S3B**). Despite the absence of significant individual DEGs with these thresholds, GSEA identified modest, coordinated pathway-level responses to VWR (**Fig. 3A**). The top 10 positively and negatively enriched gene sets from each domain are shown (**Fig. 3A; Fig. S3C–D**), with complete GSEA results provided in **Supplementary Table 1**.

**Figure 3.**
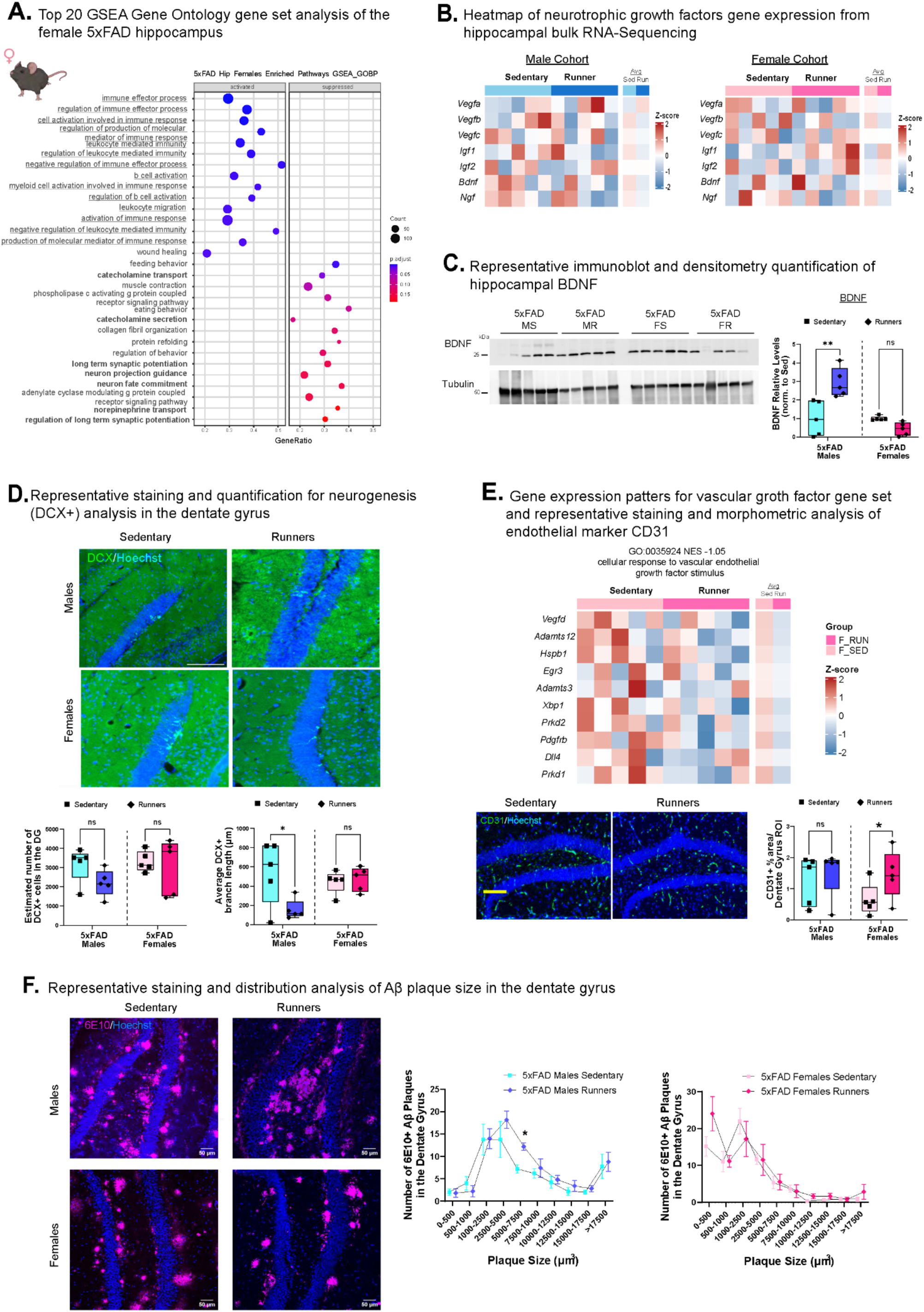
Long-term voluntary wheel running elicits modest neurotrophic, neurogenic, vascular, and amyloid-associated responses in the female 5xFAD hippocampus. **A.** Gene Set Enrichment Analysis (GSEA) of Gene Ontology Biological Process (GO:BP) terms comparing hippocampi from sedentary and VWR female 5xFAD mice. The 10 most positively and negatively enriched gene sets are shown; complete enrichment results are provided in Supplementary Table 1. **B.** Normalized hippocampal expression of neurotrophic and growth factor transcripts in sedentary and VWR male and female cohorts. **C.** Representative immunoblots and quantification of hippocampal BDNF protein abundance in male and female cohorts. **D.** Representative doublecortin (DCX) immunofluorescence in the dentate gyrus and quantification of DCX-positive immature neurons and neurite morphology in male and female cohorts. Scale bars, 50 μm. **E.** Heatmap showing normalized expression of genes (z score) comprising the cellular response to vascular endothelial growth factor stimulus GO:BP gene set in the female hippocampus, together with representative CD31 immunofluorescence and quantification of CD31-positive vascular area in the dentate gyrus of male and female cohorts. Columns in the heatmap represent individual animals followed by group- averaged expression for sedentary and VWR groups. **F.** Representative 6E10 immunofluorescence and quantification of Aβ plaque burden in the dentate gyrus of sedentary and VWR female 5xFAD mice. Plaque- size distributions were analyzed using the Kolmogorov–Smirnov (K–S) test. Unless otherwise noted, data are presented as mean ± SEM, with each point representing one animal. n = 5-6 animals/group. n.s., not significant; *P < 0.05, **P < 0.01, unpaired two-tailed Student’s t-tests.

Overall, VWR elicited a modest pathway-level transcriptional response in the female hippocampus, with most enriched GO:BP terms reaching only borderline levels of statistical significance (**Fig. 3A**). Within this limited response, VWR positively enriched several immune-related processes, including *leukocyte-mediated cytotoxicity*, *regulation of immune effector processes*, *response to interferon-γ*, and *regulation of myeloid leukocyte differentiation* (**Fig. 3A**, underlines). Conversely, VWR negatively scored neuronal processes associated with *long-term synaptic potentiation, neuron projection guidance*, and *catecholamine transport and secretion* (in bold, **Fig. 3A**, padj <0.03).

The broader GO analyses supported these transcriptional patterns. VWR positively enriched Cellular Component gene sets associated with vesicle formation, including *early endosome membrane* and *phagocytic vesicle* (**Fig. S3C**, in italics), while negatively scoring neuronal structures including *postsynaptic density membrane, dendritic shaft, neuron spine, and synaptonemal structure* (**Fig. S3C**, in bold). Beyond the immune and neuronal pathways identified by GO:BP, complementary GO:CC and GO:MF analyses highlighted positive enrichment of endosomal and proteolytic processes, including *phagocytic vesicle* and *endosomal compartments* and multiple *peptidase-associated functions* (**Fig. S3C–D,** *in italics*).

Given the negative enrichment of pathways related to synaptic plasticity and neuronal function, we next examined the expression of neurotrophic and growth factors implicated in exercise-induced hippocampal plasticity in both hippocampal transcriptomic datasets, including *Bdnf, Vegf* family members, *Ntf3*, *Ngf*, and *Fgf2*. VWR produced variable patterns in the normalized expression of neurotrophic and growth factor transcripts in either sex examined, with some genes trending higher or lower relative to sedentary animals; however, VWR did not significantly alter any individual transcript examined (**Fig. 3B**). Because BDNF plays a central role in exercise-induced hippocampal plasticity ^12^, we next quantified BDNF protein abundance by immunoblotting. Whereas males showed an exercise-induced increase in hippocampal BDNF, VWR decreased hippocampal BDNF protein levels in our independent female cohort (**Fig. 3C**), aligning with the negative enrichment of transcriptional pathways associated with synaptic plasticity and neuronal function (**Fig. 3A**).

Given the established role of BDNF in promoting adult hippocampal neurogenesis ^1,12,33^, we next asked whether these changes in BDNF abundance were accompanied by altered neurogenesis. We immunostained the dentate gyrus for doublecortin (DCX), a marker of immature neurons, and quantified DCX+ cell number and neurite length. VWR did not alter DCX+ cell number or neurite length in females (**Fig. 3D**). In contrast, VWR reduced DCX+ neurite length (with trends towards decreased DCX+ cell number) in the male cohort (**Fig. 3D**) despite the observed increase in hippocampal BDNF (**Fig. 3C**).

Vascular remodeling represents another well-established hippocampal adaptation to exercise and closely associates with adult neurogenesis ^39^. Consistent with vascular pathways emerging from the female transcriptomic analysis, VWR negatively enriched *cellular response to vascular endothelial growth factor stimulus*, with coordinated changes in genes comprising this GO term, including *Vegfd*, *Pdgfrb*, and several downstream signaling mediators (**Fig. 3E**). We therefore asked whether this transcriptional signature corresponded with structural changes in the hippocampal vasculature. Using CD31 as an endothelial marker, we quantified vascular area within the dentate gyrus and found that VWR increased CD31-positive area in females (suggesting vascular remodeling within the dentate gyrus), despite the negative enrichment of VEGF-responsive transcripts. VWR did not alter CD31-positive area in the independently studied male cohort (**Fig.3E**).

Because Aβ pathology can disrupt neuronal plasticity and cellular pathways governing protein trafficking and degradation, we further examined the endolysosomal signatures emerging from the female GSEA. In addition to enrichment of endosomal compartments and proteolytic functions (**Fig. S3C–D**, in italics), VWR positively enriched the GSEA GO BP term *lysosomal transport*, accompanied by coordinated expression changes among genes comprising this pathway (**Fig. S3E**). The convergence of endosomal, lysosomal, and proteolytic processes prompted us to examine whether these transcriptional changes coincided with alterations in hippocampal Aβ pathology following VWR. We therefore assessed hippocampal Aβ plaque burden by immunofluorescence, quantifying 6E10^+^ plaque number, average size, and size distribution in the dentate gyrus of our cohorts. VWR did not alter hippocampal plaque number, average plaque size, or plaque-size distribution in females (**Fig. 3F; Fig. S3F,G**). In contrast, VWR shifted the plaque-size distribution in males toward larger plaques, with runners exhibiting a greater proportion of plaques >2,500 µm³ and fewer plaques <1,000 µm³ relative to sedentary males (**Fig. 3F**). Together, these targeted analyses show that the molecular and cellular responses associated with female-enriched transcriptional pathways were not uniformly recapitulated in the independently studied male cohort, further demonstrating that VWR engaged overlapping but non-identical biological responses across the two cohorts.

### Long-term VWR produces limited behavioral benefits in 5xFAD mice

To determine whether the molecular responses to long-term VWR translated into functional improvements, we assessed spatial learning and memory, locomotor and anxiety-like behavior, and species-typical nesting behavior in the independently studied male and female 5xFAD cohorts. We first evaluated hippocampal-dependent spatial learning and memory using the Barnes maze. Across training days 1–4, sedentary 5xFAD males progressively reduced their latency to locate the escape hole, consistent with acquisition of the spatial task, whereas male 5xFAD runners exhibited slower task acquisition (**Fig. 4A**). Male runners also showed increased escape latency during the probe trial (**Fig. 4B**), although VWR did not alter the number of errors or time spent in the correct quadrant (**Fig. 4C-D**). Notably, male 5xFAD runners spent more time immobile during the Barnes maze (**Fig. 4E**), while VWR did not alter path efficiency or total path length (**Fig. S4B-C**). On the other hand, female 5xFAD runners similarly spent more time immobile during the Barnes maze, but VWR did not alter escape latency, number of errors, time in the correct quadrant, or total path length in the female cohort (**Fig. 4A-E; Fig. S4A-C**). Together, these findings indicate that VWR did not improve Barnes maze performance in either cohort, and that the increased escape latency in males coincided with altered task-related activity in both sexes rather than consistent deficits across spatial performance measures.

**Figure 4.**
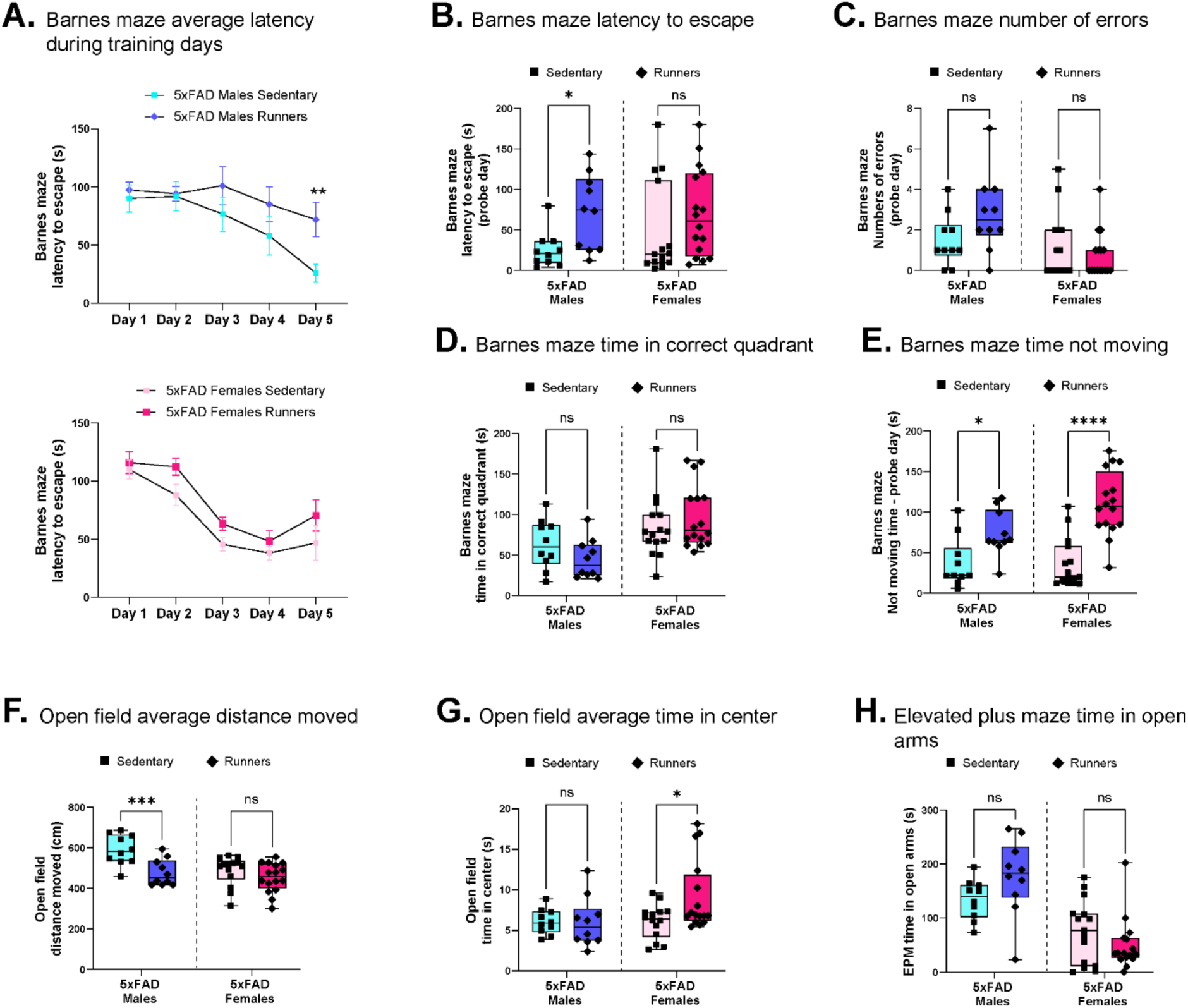
Long-term voluntary wheel running produces limited behavioral benefit in 5xFAD mice. **A.** Barnes maze escape latency across four training days in sedentary and VWR male and female 5xFAD mice. **B.** Escape latency during the Barnes maze probe trial. **C.** Number of errors during the probe trial. **D.** Time spent in the target quadrant during the probe trial. **E.** Immobility during Barnes maze testing. **F.** Total distance traveled in the open field. **G.** Time spent in the center of the open field. **H.** Time spent in the open arms of the elevated plus maze. Data are presented as mean ± SEM, with each point representing one animal except for longitudinal Barnes maze data. n = 10-16 animals/group. Barnes maze acquisition across training days was analyzed using two-way repeated-measures ANOVA. All other comparisons between sedentary and VWR groups were performed within each sex cohort using unpaired two-tailed Student’s t-tests. n.s., or lack of annotation, indicates the test was not significant; *P < 0.05.

We next used the open-field and elevated plus maze (EPM) to assess locomotor and anxiety-like behavior. In agreement with the increased immobility observed during the Barnes maze, male 5xFAD runners traveled less distance and moved at a lower average velocity in the open field than age-matched sedentary males (**Fig. 4F; Fig. S4D**). VWR did not alter open-field distance or velocity in females (**Fig. 4F; Fig. S4D**); however, female runners spent more time in the center of the open field (**Fig. 4G**), consistent with reduced anxiety-like behavior in this assay. VWR did not alter time spent in the open arms of the EPM or the open-arm entry ratio in either cohort (**Fig. 4H; Fig. S4E**). Finally, we assessed nesting as a measure of species-typical home-cage behavior. VWR did not alter nesting scores in either the male or female cohort (**Fig. S4F**).

Together, these behavioral analyses demonstrate that long-term VWR produced limited functional benefit in 5xFAD mice. It reduced locomotor activity in 5xFAD males, whereas 5xFAD females showed increased center exploration consistent with reduced anxiety-like behavior in the open field but no corresponding changes in EPM performance.

### VWR induces metabolic and transcriptional remodeling of iWAT in 5xFAD mice

Given the limited hippocampal and cognitive behavioral effects of VWR, we next examined transcriptional adaptations to VWR in inguinal white adipose tissue (iWAT), an exercise-responsive metabolic tissue that integrates energy metabolism, thermogenic adaptation, and local immune regulation ^30^. Consistent with an exercise-responsive phenotype, VWR reduced iWAT mass in the 5xFAD female cohort but did not alter iWAT mass relative to body weight in 5xFAD males (**Fig. 1G**). To determine whether VWR induced established exercise-associated metabolic adaptations in iWAT, we quantified the expression of key genes modulating adipocyte metabolic health, browning/thermogenesis, and lipogenesis by qRT-PCR (**Fig. 5A–C**). Among markers of adipocyte metabolic health, VWR increased *Adipoq* expression in females but not males, while *Ppara* remained unchanged in both cohorts (**Fig. 5A**). Assessment of the browning/thermogenic program revealed increased *Ucp1* expression in both cohorts and increased *Cidea* in females, whereas *Pgc1a* remained unchanged (**Fig. 5B**). In contrast, VWR did not alter expression of the lipogenic genes *Fasn*, *Srebf1*, or *Scd1* in either cohort (**Fig. 5C**).

**Figure 5.**
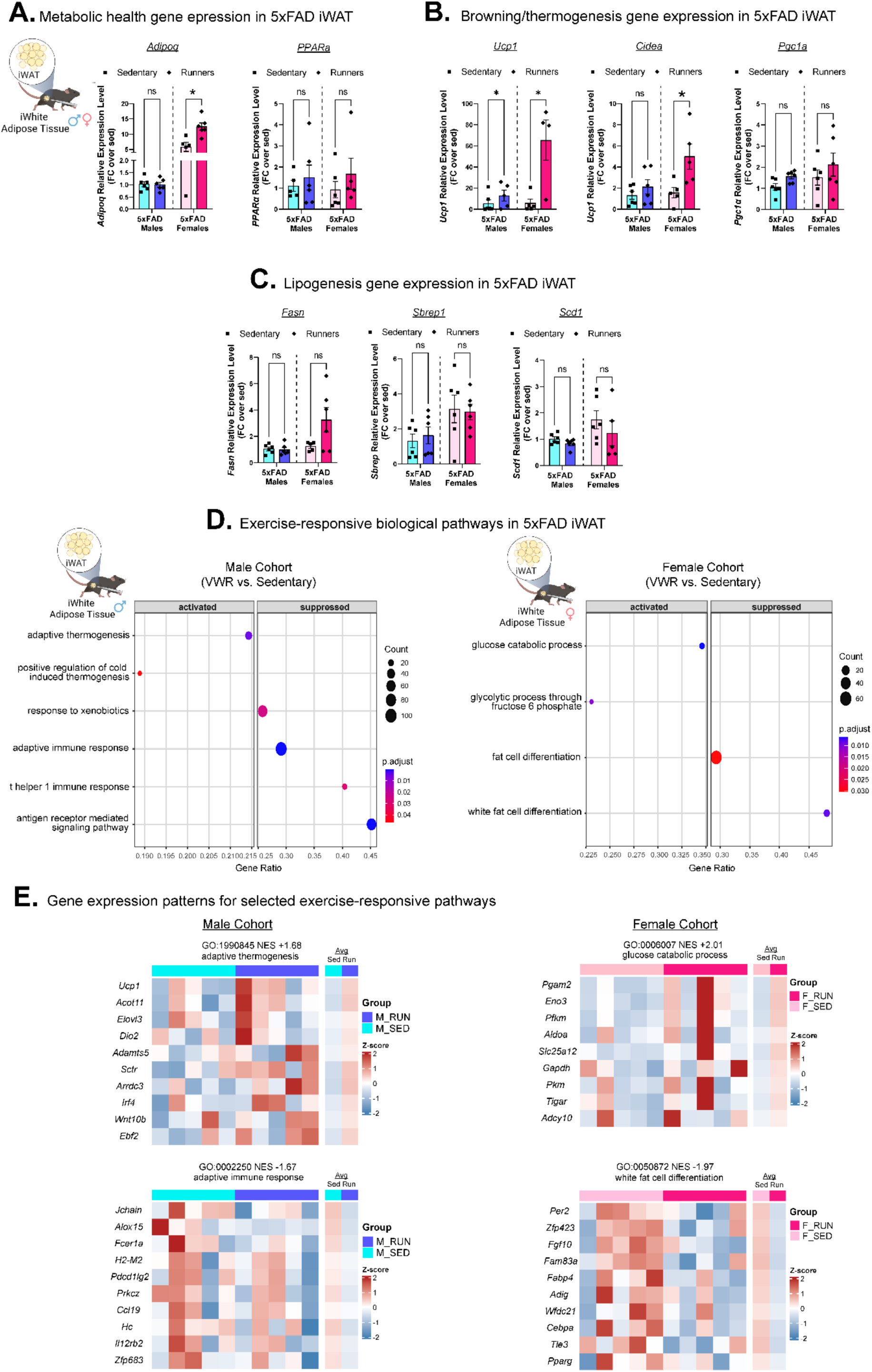
Long-term voluntary wheel running elicits metabolic and transcriptional remodeling of iWAT in 5xFAD mice. **A–C.** qRT-PCR analysis of genes associated with **A.** adipocyte metabolic health, **B.** browning and thermogenesis, and **C.** lipogenesis in inguinal white adipose tissue (iWAT) from sedentary and VWR male and female 5xFAD mice. **D.** Gene Set Enrichment Analysis (GSEA) of selected Gene Ontology Biological Process (GO:BP) terms in iWAT following VWR. Shown are thermogenic and adaptive immune-related pathways in males and glucose catabolic and adipocyte differentiation pathways in females; complete enrichment results are provided in Supplementary Tables 3. **E.** Heatmaps showing normalized expression of genes comprising the adaptive thermogenesis and adaptive immune response GO:BP gene sets in males, and the glucose catabolic process and white fat cell differentiation gene sets in females. Columns represent individual animals followed by group-averaged expression for sedentary and VWR groups. Heatmaps show VST-normalized expression as gene-wise z-scores. Unless otherwise noted, data are presented as mean ± SEM, with each point representing one animal. n = 6 animals/group for qRT-PCR and n = 5/group for RNA-seq. n.s., not significant; *P < 0.05, unpaired two-tailed Student’s t-tests.

To determine whether this targeted metabolic analysis reflected broader transcriptional remodeling, we performed bulk RNA-seq on iWAT from sedentary and runner mice in both cohorts (n=5/group). Concurrent sequencing of iWAT from both cohorts allowed direct visualization of global transcriptional variation across all samples by PCA. Male and female samples separated primarily along PC2, whereas sedentary and VWR samples showed substantial overlap within each cohort (PC1) (**Fig. S5A**), similar to what we observed in our hippocampal analysis. Given the different ages of the two cohorts, we restricted subsequent analyses to VWR versus age-matched sedentary controls within each cohort. Consistent with the limited separation observed by PCA, differential expression analysis identified no significant DEGs in male 5xFAD iWAT after multiple-testing correction (FDR < 0.10 and |log2(fold change)| ≥ 0.58), **Fig. S5B**). The 5xFAD female cohort similarly showed limited differential expression, with only a few transcripts meeting the predefined significance and fold-change thresholds: *Mfsd2a*, *Prag1*, and *Epn1* decreased following VWR, whereas *Rasd2*, and *Pygm* increased (**Fig. S5B**).

Despite the limited global separation by VWR, GSEA identified coordinated exercise-responsive pathways within each cohort (**Fig. 5D, E; Fig. S5C, D**). In males, GSEA identified 192 Biological Process, 18 Cellular Component, and 192 Molecular Function gene sets that met the significance threshold (FDR <0.05). For females, GSEA identified 166 Biological Process, 42 Cellular Component, and 166 Molecular Function gene sets that met the same FDR threshold (**Supplementary Table 2**). Both cohorts showed enrichment of several contractile/cytoskeletal and muscle-associated gene sets, which we did not interpret further as their biological relevance to iWAT remains unclear (**Fig S5C, D**). We therefore focused on pathways with established roles in adipose tissue physiology and exercise adaptation instead.

In males, VWR positively enriched *adaptive thermogenesis* and *positive regulation of cold-induced thermogenesis* (**Fig. 5D; Fig. S5C**), recapitulating a well-established exercise-responsive program in iWAT ^29,30^. To visualize the gene-level patterns underlying selected GO terms, we plotted normalized gene expression as heatmaps for the top 10 leading edge genes in each gene set, including group-averaged expression for sedentary and VWR groups (**Fig. 5E**). These profiles revealed substantial inter-individual variability in the transcriptional response to VWR, with some runners showing more pronounced shifts in pathway-associated genes than others. At the group level, however, VWR shifted the adaptive thermogenesis gene set toward higher normalized expression (**Fig. 5E**). In parallel with the thermogenic response, VWR suppressed transcriptional programs associated with adaptive immunity in male 5xFAD iWAT, including *T-helper 1 immune responses* and *antigen receptor-mediated signaling* (**Fig. 5D; Fig. S5C**). Despite variability among individual animals, the group-averaged expression profile showed an overall reduction in genes comprising the *adaptive immune response* gene set (**Fig. 5E**). Together, these findings identify a heterogeneous but coordinated exercise-responsive transcriptional program in male iWAT characterized by enhanced thermogenic signaling and suppression of adaptive immune-related pathways.

In 5xFAD females, VWR positively enriched glucose catabolism and glycolytic processes in iWAT (**Fig. 5D; Fig. S5D**). As observed in males, the expression profiles revealed substantial inter-individual variability in the response to VWR. At the group level, genes comprising the *glucose catabolic process* showed higher normalized expression in runners, consistent with the positive enrichment identified by GSEA (**Fig. 5E**). Conversely, VWR negatively enriched *fat cell differentiation* and *white fat cell differentiation,* with the group-averaged expression profile showing an overall reduction in genes comprising the white fat cell differentiation gene set (**Fig. 5D–E; Fig. S5D**). These findings identify a heterogeneous but coordinated transcriptional response in female iWAT characterized by enhanced glucose catabolism and reduced adipocyte differentiation, distinct from the thermogenic and immune-related programs identified in the independently studied male cohort. Furthermore, these peripheral adaptations provide independent evidence of a physiologically effective exercise stimulus despite the limited CNS and behavioral benefits of VWR.

## DISCUSSION

In this study, we investigated the effects of long-term voluntary wheel running on behavioral, neuropathological, cellular, and molecular outcomes in 5xFAD mice. Both cohorts sustained voluntary running throughout the intervention and exhibited exercise-responsive transcriptional remodeling of iWAT. In the hippocampus, VWR produced modest, pathway-level transcriptional responses accompanied by changes in selected synaptic, mitochondrial, neurotrophic, and monoaminergic proteins. Despite this molecular responsiveness, VWR produced little evidence of broad disease modification: exercise did not reduce hippocampal Aβ pathology, increase adult neurogenesis, or improve spatial behavioral performance, and transcriptional changes in neuroimmune pathways were not accompanied by detectable changes in the measured cellular endpoints. Several exercise-responsive molecular adaptations likewise occurred without corresponding structural or functional changes. Together, these findings distinguish engagement of exercise-responsive biology from broad modification of AD-like pathology and suggest that the rapidly progressive 5xFAD disease environment may limit the translation of exercise-induced molecular responses into neuroprotective outcomes.

The limited disease-modifying effects of VWR observed here are consistent with previous evidence that voluntary exercise does not uniformly ameliorate the 5xFAD phenotype. Indeed, voluntary exercise has produced markedly heterogeneous outcomes in 5xFAD mice, ranging from improved cognition, neurogenesis, neurotrophic signaling, and amyloid pathology ^11,12^ to minimal or no CNS benefit ^17,19^. Such variability is also not unique to the 5xFAD transgenic model; exercise studies in APP/PS1 mice similarly report responses that vary with disease stage and experimental paradigm, ranging from broad cognitive and neuropathological improvement ^14^ to more restricted or absent effects ^40^. This heterogeneity likely reflects, among other things, the strong context dependence of exercise efficacy, including differences in disease stage and severity, intervention timing, exercise modality, and cumulative exercise exposure. These factors may be particularly consequential in 5xFAD mice, where rapidly progressing amyloid pathology and associated neural dysfunction could increasingly constrain the capacity of exercise-induced adaptations to modify the underlying disease process.

In agreement with this, the magnitude of hippocampal transcriptional remodeling observed here was considerably more modest than the extensive exercise-responsive transcriptional changes reported in the healthy brain ^9,15,16,41^. In our hands, VWR produced predominantly pathway-level shifts, with limited differential expression at the individual-gene level. This modest transcriptional response may suggest that the 5xFAD disease environment constrains the molecular plasticity normally elicited by exercise. Consistent with this interpretation, voluntary running also produced minimal global transcriptional remodeling in male 5xFAD mice in a previous study, despite eliciting an extensive transcriptional response in WT animals ^19^. Alternatively, the magnitude of the transcriptional response may reflect the exercise stimulus itself. VWR permits self-selected activity, resulting in substantial individual variation in exercise intensity and cumulative workload. The exercise burden achieved here may consequently have been insufficient and/or too variable to elicit broader hippocampal remodeling.

One feature of the hippocampal response to VWR was that transcriptional adaptations did not consistently translate into physiological and functional changes typically associated with exercise. For example, VWR altered several pre- and postsynaptic proteins, yet excitatory and inhibitory synapse density remained unchanged. This divergence may reflect remodeling of existing synapses rather than changes in synapse number, as alterations in presynaptic vesicle organization and trafficking or postsynaptic assembly can modify synaptic composition and function without requiring structural gain or loss. Neurotrophic, mitochondrial, and vascular responses were similarly dissociated. Rather than engaging a coordinated neurotrophic–neurogenic–vascular program, VWR activated different components of this canonical exercise response independently in the 5xFAD hippocampus. For example, in males, increased BDNF failed to support neurogenesis, potentially reflecting reduced progenitor competence or an inhospitable neurogenic niche in the setting of Aβ pathology ^42^. In females, vascular remodeling occurred despite reduced BDNF and unchanged neurogenesis, suggesting that the vascular response can proceed independently of neurogenic adaptation. Together, these findings suggest that established amyloid pathology disrupts the coupling among neurotrophic signaling, neurogenesis, and vascular remodeling that normally characterizes hippocampal adaptation to exercise. Notably, these transcriptional and molecular adaptations to VWR emerged despite unchanged Aβ pathology, suggesting that exercise-responsive neural plasticity can persist independently of measurable amyloid clearance. Similar dissociations between exercise-induced adaptations and plaque burden have been previously reported in 5xFAD mice and other models of Aβ neurotoxicity ^11,17,43^.

This limited translation extended to behavioral outcomes. VWR did not improve hippocampal-dependent cognitive performance, despite extensive evidence that exercise enhances spatial learning and memory in both aged and AD mouse models. The unexpected impairment in Barnes maze acquisition coincided with reduced spontaneous locomotion and greater task-related immobility, complicating its interpretation as a selective deficit in spatial learning, as altered motivation or task engagement may also have contributed to this phenotype. This consideration is particularly relevant in 5xFAD mice, which exhibit abnormal exploratory and avoidance behaviors that complicate conventional interpretation of cognitive and anxiety-related assays ^27^. The accompanying remodeling of monoaminergic signaling in males provides another potential contributor to this behavioral phenotype. Serotonergic signaling, particularly through 5-HT1A receptors, plays a well-established but circuit-dependent role in anxiety, exploratory behavior, and behavioral responses to stress ^44^. VWR increased hippocampal 5-HT1A and decreased DRD4 abundance in males, potentially connecting alterations in monoaminergic signaling to changes in exploratory drive or task engagement ^45^. However, the functional consequences of these receptor changes cannot be inferred from total hippocampal abundance alone. The broader VWR literature further illustrates this behavioral complexity, with voluntary running producing anxiolytic ^46^, anxiogenic ^47^, or discordant exploratory phenotypes depending on the behavioral paradigm and experimental context, including in 5xFAD mice ^17^. These behavioral data underscore the importance of distinguishing cognitive performance from exercise-induced changes in locomotion, exploration, and affective state. Together, these findings suggest that the 5xFAD hippocampus retains the capacity to engage individual components of the exercise response, while established disease may limit its ability to drive coordinated cellular and functional neuroprotective adaptations typically associated with exercise.

Importantly, the limited CNS response in our study did not reflect a generalized failure to adapt to VWR. There is substantial evidence that subcutaneous adipose tissue (like iWAT) is better adapted for metabolically favorable lipid storage and energy utilization, while visceral adipose tissue is more closely associated with immune signaling and inflammatory regulation. Thus, through an unbiased approach, iWAT retained transcriptional exercise responsiveness in 5xFAD runners, providing an important peripheral contrast to the comparatively restricted hippocampal response. In 5xFAD males, VWR engaged a canonical thermogenic program in iWAT ^29,30^, while suppressing multiple adaptive immune-related transcriptional programs instead. In females, VWR instead favored glucose catabolism while suppressing adipocyte differentiation, accompanied by a reduction in iWAT mass consistent with broader remodeling of adipose metabolism and energy storage. The preferential enrichment of thermogenic pathways in 5xFAD male iWAT parallels previous findings that VWR induced robust thermogenic remodeling of iWAT in non-transgenic male, but not female, mice despite greater running distances in females ^29^. Our findings demonstrate that successful peripheral exercise adaptation can coexist with comparatively limited CNS disease modification.

An important consideration in interpreting the cohort-specific responses is that we intentionally staggered the age at VWR initiation to account for the accelerated progression of amyloid pathology in female 5xFAD mice ^27,28^. Females began VWR at 3 months of age and males at 6 months, with both cohorts undergoing the same 16-week intervention. This design aimed to better align amyloid disease burden across cohorts rather than chronological age. At endpoint, sedentary males and females showed comparable hippocampal plaque number and glial morphometrics, although total plaque volume remained lower in females, indicating that the staggered design reduced but did not eliminate differences in disease-associated pathology between cohorts. Nevertheless, chronological age remained intrinsically linked to cohort, precluding attribution of the distinct molecular and physiological responses observed here specifically to biological sex. These differences are nonetheless notable given the well-established influence of sex on AD pathophysiology ^48^ and growing evidence that biological sex also shapes molecular and physiological adaptations to exercise ^8,9^. Thus, although our design does not permit a formal assessment of sex-dependent exercise responses, the distinct patterns observed across the independently studied male and female cohorts reinforce the importance of incorporating biological sex into studies examining exercise responsiveness in AD. Future studies that independently manipulate age, disease burden, and sex will be necessary to determine how each contributes to the heterogeneity of exercise-induced neuroprotection.

Collectively, our findings distinguish responsiveness to exercise from disease modification in the 5xFAD model. Long-term VWR elicited clear exercise-responsive remodeling of iWAT and modest, coordinated molecular responses within the hippocampus, yet produced little improvement in established neuropathology or behavioral function. Moreover, several canonical components of the hippocampal exercise response failed to converge into the coordinated cellular adaptations associated with neuroprotection. Together, these findings suggest that the 5xFAD brain remains responsive to exercise but may be unable to mount a molecular and cellular response of sufficient magnitude or coordination to substantially modify disease-related outcomes. In rapidly progressive neurodegenerative disease, the biological engagement of exercise may therefore be insufficient to ensure disease modification, underscoring the importance of disease stage, exercise burden, and biological context when designing and interpreting exercise-based interventions.

## METHODS

### Animals

For this work, we utilized heterozygous 5xFAD transgenic mice of both sexes (MMRRC Strain: 034840- JAX | B6SJL Tg(APPSwFlLon,PSEN1*M146L*L286V)6799Vas/ Mmjax), generated by in-house breeding of 5xFAD+ and C57BL/6 breeder pairs. 5xFAD+ individuals (littermates) were randomly allocated to either the sedentary or runner groups. This line has been backcrossed and maintained on a C57BL/6 background for more than ten generations in our laboratory. Animals were housed in standard shoebox-style cages with filter tops under a 12:12 light-dark cycle. Mice were not fasted before tissue collection, and they were allowed free access to standard NIH-07 mouse chow throughout the experiment. We did not track their food consumption during this study. All animal experimentation adhered to NIH guidelines and was performed by blinded investigators (when possible) and was approved by and performed in accordance with the University of Southern California (USC) Institutional Animal Care and Use Committee.

### Voluntary running exercise protocol

To equilibrate the influence of single housing on animal behavior, each mouse from sedentary/runner cohorts was single-caged in standard vivarium polypropylene cages (290 × 180 × 160 mm) throughout the experiment. Bedding for all cages was changed to sawdust and included two nestlets and manzanita sticks for enrichment. Each running mouse had free access to a running wheel (10.16 cm in diameter) connected to a counter to record the running distance (Columbus Instruments monitoring software). The in-cage monitoring system recorded the number and timing of wheel revolutions at 1-min intervals. Sedentary controls were also given free access to a locked wheel to control for enrichment. The study was initiated at 3 months of age in females and 6 months of age in males, and continued until euthanasia and tissue collection at approximately 7 months of age in females and 12 months of age in males. Investigators monitored animal welfare, physical activity, and running performance daily. Mice retained access to their assigned free or locked wheels throughout behavioral testing, for a total intervention of approximately 16 weeks and 5 days. Wheel access was terminated on the morning following the final neurocognitive assessment.

### Tissue collections

Tissues were collected between 9:00 and 11:00 AM (ZT3–ZT5; lights on at 6:00 AM), several hours after the preceding dark-phase running period. Body weight was measured before euthanasia and tissue collection. We collected tissues as described previously with minor modifications ^49,50^. Briefly, animals were anesthetized with 3.8% Avertin solution prepared from 2,2,2-tribromoethanol (Sigma, T48402) and 2-methyl-2-butanol (Sigma, 152463) in Milli-Q water. Euthanasia was completed via diaphragm transection followed by transcardial perfusion with 60 mL of ice-cold 1x PBS. We microdissected the left hemibrains into cortex and hippocampus and flash-froze these regions in liquid nitrogen for subsequent protein analyses. We post-fixed the right hemibrains in 4% paraformaldehyde (Fisher Scientific, 50-980-495) for less than 24 hours, transferred them to 30% (w/v) sucrose for 24 hours, embedded them in OCT compound (Tissue-Tek, 4583), and froze them in 2-methylbutane (VWR, 103525-278) vapor phase cooled in liquid nitrogen. Inguinal white adipose tissue (iWAT) was flash-frozen in liquid nitrogen and/or embedded in OCT following dissection, and stored at −80°C until used.

### Protein Isolation

We isolated protein lysates from hippocampal tissue using mechanical homogenization through sequential needle extrusion in RIPA buffer supplemented with 1X Halt protease and phosphatase inhibitor cocktail. Briefly, we extruded flash-frozen hippocampi through 1 mL syringes containing 250 μL of ice-cold RIPA lysis and extraction buffer (Thermo Fisher Scientific, 89900) supplemented with 1X Halt protease and phosphatase inhibitor cocktail (Thermo Fisher Scientific, 78430). We homogenized the tissue by repeatedly passing the suspension first through an 18-gauge needle (Thermo Fisher Scientific, 14-826A) and then through a 23-gauge needle (Thermo Fisher Scientific, 14-826-5G) until no visible fragments remained. We clarified the hippocampal lysates by centrifugation at 8,000 × g at 4°C for 10 minutes and collected the supernatant as the crude protein lysate. We then diluted the supernatant 1:1 (v/v) with fresh ice-cold RIPA buffer supplemented with 1X Halt protease and phosphatase inhibitor cocktail to generate working lysates. We quantified protein concentrations using the Pierce BCA Protein Assay Kit (Thermo Fisher Scientific, 23227) and stored lysates at −80°C until analysis.

### Immunoblot analyses

For immunoblotting of hippocampal lysates, we prepared 20 µg of homogenized protein per sample with NuPAGE LDS sample buffer (Thermo Fisher Scientific, NP0007) and NuPAGE sample reducing agent (Thermo Fisher Scientific, NP0004) as described previously ^50^, and separated proteins on 4–15% Criterion TGX Precast Midi Protein Gels (Bio-Rad, 5671085). We transferred proteins onto methanol activated 0.45 µm PVDF membranes (Thermo Fisher Scientific, 88518) and blocked the membranes in 5% nonfat dry milk in 1× TBST (1× TBS with 0.05% Tween-20) at room temperature for 1 hour. We incubated membranes overnight at 4°C with primary antibodies against BDNF, COX10, NDUFS1, PSD95, synapsin-1, synaptophysin, UQCRFS1, DRD4, 5-HT1A, typically at a 1:1000 dilution (**Supplementary Table S3**). Corresponding secondary antibodies were diluted at 1:5000. After primary incubation, we incubated membranes with secondary antibodies for 1 hour at room temperature with gentle shaking. We imaged membranes using a ChemiDoc imaging system (Bio-Rad, 12003153) and performed densitometry analyses using ImageJ analysis software, normalizing densitometry signals to loading controls and expressing values relative to the average of sedentary individuals by sex.

### Immunofluorescence staining and imaging

Hemibrains were embedded in OCT (Tissue-Tek, 4583), frozen in 2-methylbutane (VWR, 103525-278) cooled in liquid nitrogen, and stored at −80°C until sectioning. Brains were sectioned at 20 μm on a Leica cryostat and mounted onto SuperFrost Plus slides (VWR, 48311-703). Sections were permeabilized in 0.1% Triton X-100 for 15 min and blocked in 4% BSA for 1 h. Brain sections were incubated with primary antibodies overnight at 4°C, followed by the appropriate secondary antibodies for 1 h at room temperature (**Supplementary Table 3**). Sections were counterstained with Hoechst (Thermo Fisher Scientific, 62249; 1:5000) and mounted with ProLong Glass Antifade Mountant (Invitrogen, P36984). Sections were washed three times for 5 min in 1× PBS between each step. For sections stained for DCX, vGLUT1, vGAT, or Gephyrin, we performed antigen retrieval before permeabilization by heating sections in 1× citrate buffer (MilliporeSigma, 21545) under high pressure for 6 min in a pressure cooker.

Brain sections were imaged on a Stellaris 5 confocal microscope (Leica Microsystems, 8119637) at 20× or 63× magnification, acquiring 10-μm z-stacks at 1024 × 1024 pixels. When appropriate, sections were imaged with an ECHO Revolution epifluorescent microscope at 10x magnification, and tile scans of entire sections were acquired stacks with Z-stacks were collapsed into maximum intensity projections, yielding 2D images which were used for analysis.

### Immunostaining quantification

We used Imaris imaging software (Oxford Instruments, version 10.2) for plaque and glial analyses.

Volumetric surfaces of amyloid-β plaques, astrocytes, and microglia were generated based on immunofluorescence signal intensity. For astrocytes and microglia, surfaces selected were filtered by size to include positive structures above a 10µm^3^ threshold. For plaques and glia, average volumes per ROI were calculated from 3 replicates per animal.

Quantification of doublecortin-positive cells was performed in FIJI using the Neuroanatomy Simple Neurite Tracer (SNT v5.0.5) plugin to manually label DCX+ cells and trace neurite branches ^51^. Three hippocampal brain sections per animal were analyzed and averaged to generate total branch length sums for each individual. This was defined as the sum of all neurite paths traced and was calculated in Excel from average branch length sums across replicates for each individual. Estimates for the number of total DCX+ cells per individual were determined by design-based stereology for standardized quantification of adult neurogenesis ^52^, with minor modifications. Briefly, we used raw counts from 2-dimensional representative brain sections from each individual to estimate 3-dimensional populations in the entire hippocampus. We estimated the rostrocaudal extent of the adult mouse hippocampus as approximately 2.4 mm, corresponding to ∼120 sections at 20 μm thickness, based on the Allen Mouse Brain Atlas. After establishing our size parameters, we quantified every 16th section of tissue for every individual to cover the proximal, medial, and anterior dentate gyrus.

For quantification of synapses, the open-source FIJI plugin Synbot ^37^ was utilized to count colocalizations of presynaptic and postsynaptic markers. The average number of synapses per µm2 was calculated from 3 hippocampal brain sections, in which 3 ROIs of uniform dimensions at 63x magnification were collected from each hippocampal brain section, totaling 9 replicates per animal. These values were then divided by the ROI area, 184.70 µm^2^. Two-channel colocalization (RG) was performed on each image, wherein pre-processing of images within the Synbot interface included noise reduction with a rolling ball radius of 50 pixels and Gaussian blur of 0.570. Thresholding included 20% histogram thresholding. Whole image ROIs were selected, as images taken were of uniform dimensions. Red and green minimum and maximum pixel sizes were both set to 4 and infinity, respectively. Circular approximation analyses were performed.

### iWAT RNA extraction for bulk RNA sequencing and real-time PCR

Total RNA was extracted from inguinal white adipose tissue (iWAT) using an RNeasy Lipid Tissue Mini Kit (QIAGEN, 74804) and the manufacturer’s protocol. RNA purity and quantity were determined by spectrophotometry using a NanoDrop (Thermo Scientific, ND-ONE-W). Extracted RNA was reverse transcribed into cDNA using SuperScript IV VILO Master Mix (ThermoFisher Scientific, 11756050). Primer pairs were designed using IDT™ oligo nucleotide entry tool, using predetermined sequences confirmed by the National Center for Biotechnology Information (NCBI) (**Supplementary Table S3**).

### Real-time PCR

mRNA abundance was quantified using a QuantStudio 3 Real-Time PCR System (ThermoFisher, A28567) and Fast SYBR Green Master Mix (ThermoFisher, 4385612). Cycling conditions followed the manufacturer’s protocol. Gene expression of targets was normalized to cyclophilin as the endogenous control.

### Gene expression analysis

Total RNA was extracted from flash-frozen hippocampal tissue by Novogene (Novogene Corporation Inc., Sacramento, CA, USA), from five biological replicates per sex/genotype group. RNA quality was assessed on a Bioanalyzer at Novogene. mRNA libraries were prepared, and bulk RNA-seq was performed by Novogene as previously described ^49^. Briefly, high-throughput sequencing was carried out on a NovaSeq 6000 PE150 platform (Illumina, San Diego, CA, USA). We used HISAT2 (v2.2.0) to build the reference index and align clean reads to the reference genome. Read counts were generated using featureCounts (v1.5.0-p3).

#### Data availability

Raw sequencing data is available upon request.

#### Bioinformatics analysis

The analyses for hippocampal tissue and inguinal adipose tissue were performed in R (4.5.1) ^53^, based on previous methods. Briefly, raw gene counts were imported and filtered to retain only protein-coding genes, duplicates were collapsed by retaining the entry with the highest total count across samples. Genes were filtered for a minimum expression threshold (≥10 counts in ≥3 of 5 samples per group). Differential expression analysis was performed separately within each cohort using sedentary animals as the reference group ^54,55^. We defined differentially expressed genes as those with a |log2(FoldChange)| ≥ 0.58 and an FDR < 0.10. Volcano plots were generated using the EnhancedVolcano package with these thresholds. Enrichment analysis was performed using fgseaMultilevel with genes ranked by log2 fold change for gene ontology Biological Processes (GO:BP), Cellular Component (GO:CC), and Molecular Function (GO:MF) sets retrieved from MSigDB M5 collection via msigdbr ^56–58^. Gene set size was established at 25 to 500 genes. Significant pathways were defined by Benjamini-Hochberg adjusted p <0.05. Results were visualized in bubble plots using the ggplot2 package version 4.0.3 ^59^. Gene ratio was calculated as the number of leading-edge genes divided by the total number of genes in the corresponding pathway ^60^. Enrichment results were visualized as bubble plots showing either the top pathways ranked by adjusted p-value or targeted pathway subsets relevant to each comparison. For selected enriched GOID pathways, VST-normalized expression for the top 10 leading edge genes is shown, z-scored per sample and as group averages visualized on heatmaps using ComplexHeatmap version 2.24.1. For selected neurotrophic factors, VST-normalized expression of the selected genes is shown, z-scored per sample and as group averages visualized on heatmaps using ComplexHeatmap version 2.24.1 ^61,62^. Claude and ChatGPT5 LLMs were used for R code editing and debugging of this bioinformatics pipeline. Although transcriptomic analyses were performed independently within each cohort, we assessed selected downstream molecular and cellular endpoints in both cohorts to determine whether the exercise-responsive biology identified by transcriptomics was shared across cohorts or specific to one. All statistical analyses remained restricted to VWR versus sedentary comparisons within each sex cohort.

### Mouse phenotyping and neurocognitive behavior

All behavioral experiments listed below were performed by double-blinded investigators. Unless otherwise stated, group sizes were as follows: 5xFAD sedentary males (n=10), 5xFAD sedentary females (n=15), 5xFAD runner males (n=10), 5xFAD runner females (n=16). Group size changes across phenotyping tests reflect unrelated veterinary flags that required an individual’s removal from the study, such as fighting injuries, flooded cages, and/or prolapses.

#### Barnes Maze

Barnes maze testing was performed to assess spatial learning and memory. The maze consisted of a 75-cm-diameter circular platform containing 20 evenly spaced holes, with an escape box positioned beneath one target hole. Distinct, stationary spatial cues surrounded the maze and remained constant throughout testing. Mice received three trials per day, with each trial lasting up to 3 min. At the beginning of each trial, mice were placed in a start box at the center of the maze for 10 s before being released to explore. On day 1, mice were habituated to the maze and allowed to explore freely for 3 min under illumination from a red light. On day 2, mice underwent an initial guided training trial in which the buzzer (80 dB) and overhead light (400 lux) were activated and mice were guided to the target escape box, where they remained for 1 min before returning to their home cage. For subsequent acquisition trials, mice were released from the start box with the buzzer and light activated and allowed to explore for up to 3 min or until they entered the escape box. Days 3 and 4 followed the same acquisition procedure without the initial guided training trial. On day 5, mice underwent a probe trial in which the escape box was replaced with a decoy box and mice were allowed to explore the maze for 3 min. Behavior was recorded and analyzed using EthoVision software (Noldus, version 16).

#### Elevated Plus Maze

The maze consisted of four arms arranged in a plus configuration, with two open and two closed arms. Mice were placed in the center of the maze and allowed to explore for 5 min. Time spent in the open arms and open-arm entry ratio were quantified using EthoVision software (Noldus, version 16).

#### Open field

We performed open field studies as reported previously ^50^. Briefly, mice were placed in a 50 × 50-cm arena and allowed to explore freely for 5 min. EthoVision software (Noldus, version 16) was used to quantify total distance traveled, time spent in the center, and average velocity.

#### Nest Building

Nest building was conducted as previously described ^50^. Briefly, mice were placed in individual testing cages with one ∼3 g piece of nestlet and no other nesting materials or environmental enrichment items. After 24 h, nests were evaluated on a scale from 0 (no nestlet interaction) to 4 (fully developed nest) based on the shredding and morphology of nests built.

### Biochemical statistical analysis

All immunoblotting, immunostaining, behavioral, and biochemical data were analyzed as described in figure legends. Statistical significance was defined as p < 0.05. Unless otherwise noted, data are represented as means with standard error of the mean using GraphPad Prism 10.4.2 (La Jolla, CA). When possible, data analyses were conducted in a blinded fashion. All data were prepared for analysis with standard spreadsheet software (Microsoft Excel).

## ACKNOWLEDGEMENTS

We thank members of the lab, past and present, as well as former and current colleagues and collaborators, for their helpful contributions.

## AUTHOR CONTRIBUTIONS

K.A, K.G, K.L, T.J, E.M., A.B, M.H, J.L., E.M., J.A.G.L, K. E.Y., C.M.H, performed the experimental work and reviewed the manuscript. K.A. and C.J.C. wrote and edited the manuscript. R.V.D.K. and I. K. aided in experimental data analysis and editing of the manuscript. All authors read and approved the final manuscript. C.J.C. conceptualized the experiments outlined here and obtained (or helped obtain) all associated funding

## SUPPLEMENTARY FIGURE LEGENDS

**Supplementary Figure 1. Baseline pathological characteristics of the independently studied 5xFAD cohorts. A.** Experimental design aligned with the published 5xFAD disease-progression timeline adapted from the Alzforum 5xFAD model database. Female 5xFAD mice began sedentary locked-wheel or voluntary wheel running (VWR) exposure at 3 months of age and male 5xFAD mice at 6 months of age, with both cohorts undergoing 16 weeks of intervention. The staggered intervention onset was selected to account for the earlier progression of amyloid pathology in female 5xFAD mice and to better align amyloid disease burden across cohorts. **B.** Comparison of hippocampal Aβ plaque burden and astrocyte and microglial morphometric measures in sedentary male and female 5xFAD mice at endpoint. Despite differences in chronological age, sedentary male and female cohorts showed comparable hippocampal plaque number and astrocyte/microglial morphometric measures, while total plaque volume differed between cohorts, consistent with the study design intended to better align amyloid disease burden across cohorts. Data are presented as mean ± SEM, with each point representing one animal. n = 5-6 animals/group. Statistical comparisons were performed using a two-tailed Student’s t-test between sexes. ** P < 0.01. n.s., not significant.

**Supplementary Figure 2. Transcriptomic and glial responses to long-term voluntary wheel running in the male 5xFAD hippocampus. A.** Principal component analysis (PCA) of hippocampal RNA-seq profiles from sedentary and VWR male 5xFAD mice. **B.** Volcano plot showing differential gene expression following VWR, differential-expression thresholds as indicated. **C.** Gene Set Enrichment Analysis (GSEA) of Gene Ontology Cellular Component (GO:CC) terms comparing sedentary and VWR male hippocampi. The 15 most positively and negatively enriched gene sets are shown. **D.** GSEA of Gene Ontology Molecular Function (GO:MF) terms comparing sedentary and VWR male hippocampi. The 15 most positively and negatively enriched gene sets are shown. **E.** Quantification of GFAP-positive astrocyte number. **F.** Distribution of GFAP- positive structures across size ranges in the male dentate gyrus. GFAP-size distributions were analyzed using Kolmogorov–Smirnov (K–S) test. **G.** Representative IBA1-positive microglia classified across morphological states ranging from steady-state and hyper-ramified to amoeboid morphologies. Scale bar = 10 µm. **H.** Distribution of IBA1-positive microglial morphological states in sedentary and VWR male 5xFAD mice Complete GSEA results are provided in Supplementary Table 1. RNA-seq, n = 5/group. Histological analyses, n = 5-6/group. Data are presented as mean ± SEM where applicable, with each point representing one animal. n.s., not significant, unpaired two-tailed Student’s t-tests.

**Supplementary Figure 3. Transcriptomic, endolysosomal, and amyloid-associated responses to long-term voluntary wheel running in the female 5xFAD hippocampus**. A. Principal component analysis (PCA) of hippocampal RNA-seq profiles from sedentary and VWR female 5xFAD mice. B. Volcano plot showing differential gene expression following VWR. No individual transcripts met the differential-expression thresholds indicated. C. Gene Set Enrichment Analysis (GSEA) of Gene Ontology Cellular Component (GO:CC) terms comparing sedentary and VWR female hippocampi. The 15 most positively and negatively enriched gene sets are shown. D. GSEA of Gene Ontology Molecular Function (GO:MF) terms comparing sedentary and VWR female hippocampi. The 15 most positively and negatively enriched gene sets are shown. E. Heatmap showing normalized expression of genes comprising the positively enriched lysosomal transport GO:BP gene set. Columns represent individual animals followed by group-averaged expression for sedentary and VWR groups. **F.** Quantification of average hippocampal Aβ plaque size in sedentary and VWR female 5xFAD mice. **G.** Distribution of hippocampal Aβ plaques across size ranges in sedentary and VWR female 5xFAD mice. Complete GSEA results are provided in Supplementary Table 1. RNA-seq, n = 5/group; histological analyses, n = 5-6/group. Data are presented as mean ± SEM where applicable, with each point representing one animal. n.s., not significant, unpaired two-tailed Student’s t-tests.

**Supplementary Figure 4. Additional behavioral outcomes following long-term voluntary wheel running in 5xFAD mice. A–C.** Additional Barnes maze measures in sedentary and VWR male and female 5xFAD mice, including **A.** Barnes moving time, **B.** path efficiency, and **C.** total path length. **D.** Average velocity during the open-field test. **E.** Open-arm entry ratio during the elevated plus maze. **F.** Nest-building scores as a measure of species-typical home-cage behavior. Data are presented as mean ± SEM, with each point representing one animal. n = 10-16 animals/group. n.s., or lack of annotation, indicates the test was not significant; *P < 0.05, **P < 0.01, unpaired two-tailed Student’s t-tests within sex.

**Supplementary Figure 5. Transcriptomic responses to long-term voluntary wheel running in 5xFAD iWAT. A.** Principal component analysis (PCA) of iWAT RNA-seq profiles from sedentary and VWR male and female 5xFAD mice. **B.** Volcano plots showing differential gene expression following VWR within the male and female cohorts. No individual transcripts met the differential-expression thresholds indicated in the figure in males, whereas a limited number of transcripts met these thresholds in females. **C.** Gene Set Enrichment Analysis (GSEA) of Gene Ontology (GO) terms comparing sedentary and VWR male iWAT. The 10 most positively and negatively enriched gene sets within each GO domain are shown. **D.** GSEA of GO terms comparing sedentary and VWR female iWAT. The 10 most positively and negatively enriched gene sets within each GO domain are shown. Complete iWAT GSEA results are provided in Supplementary Table 3. RNA-seq, n = 5/group.

