## Supplementary Figures for "Long-term voluntary exercise reveals limited translation of hippocampal molecular responses into neuroprotection in 5xFAD mice"

### Supplementary Figure 1

**A.** 5xFAD transgenic line pathology timeline relative to experimental timelines

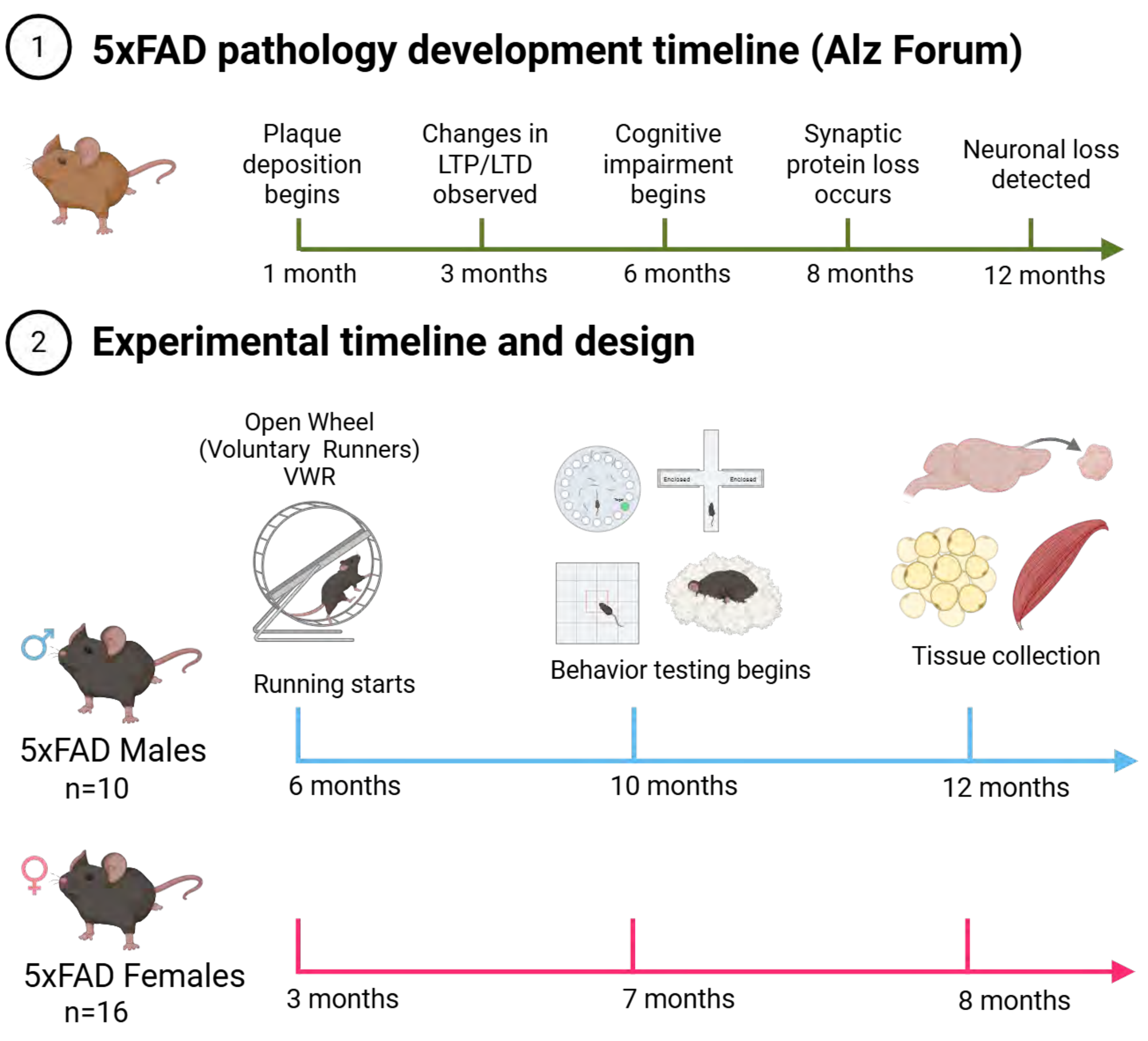


**B.** Hippocampal Aβ plaque number and volume in sedentary male and female 5xFAD cohorts

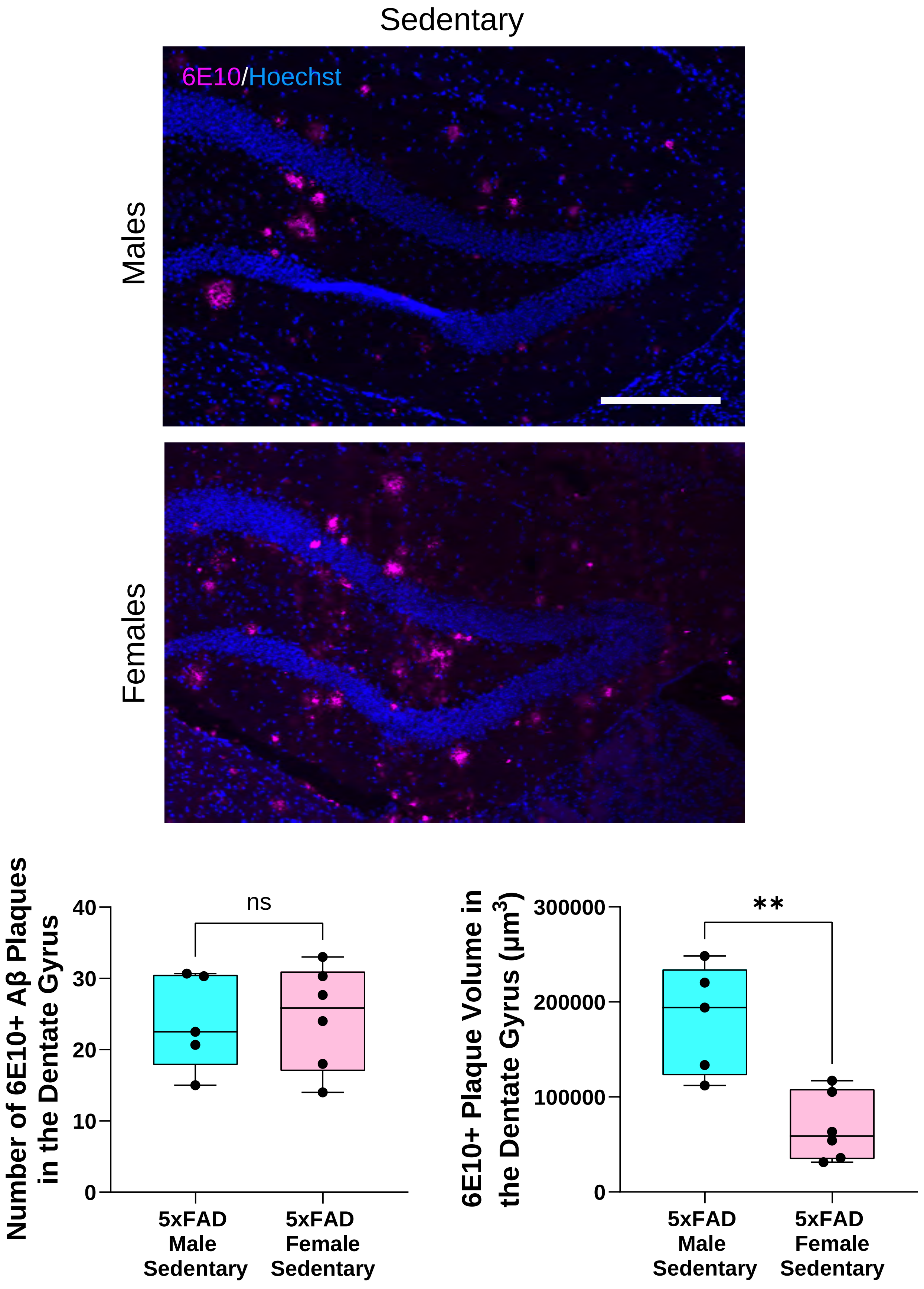


**C.** Hippocampal astrocyte and microglial morphometrics in sedentary male and female 5xFAD cohorts

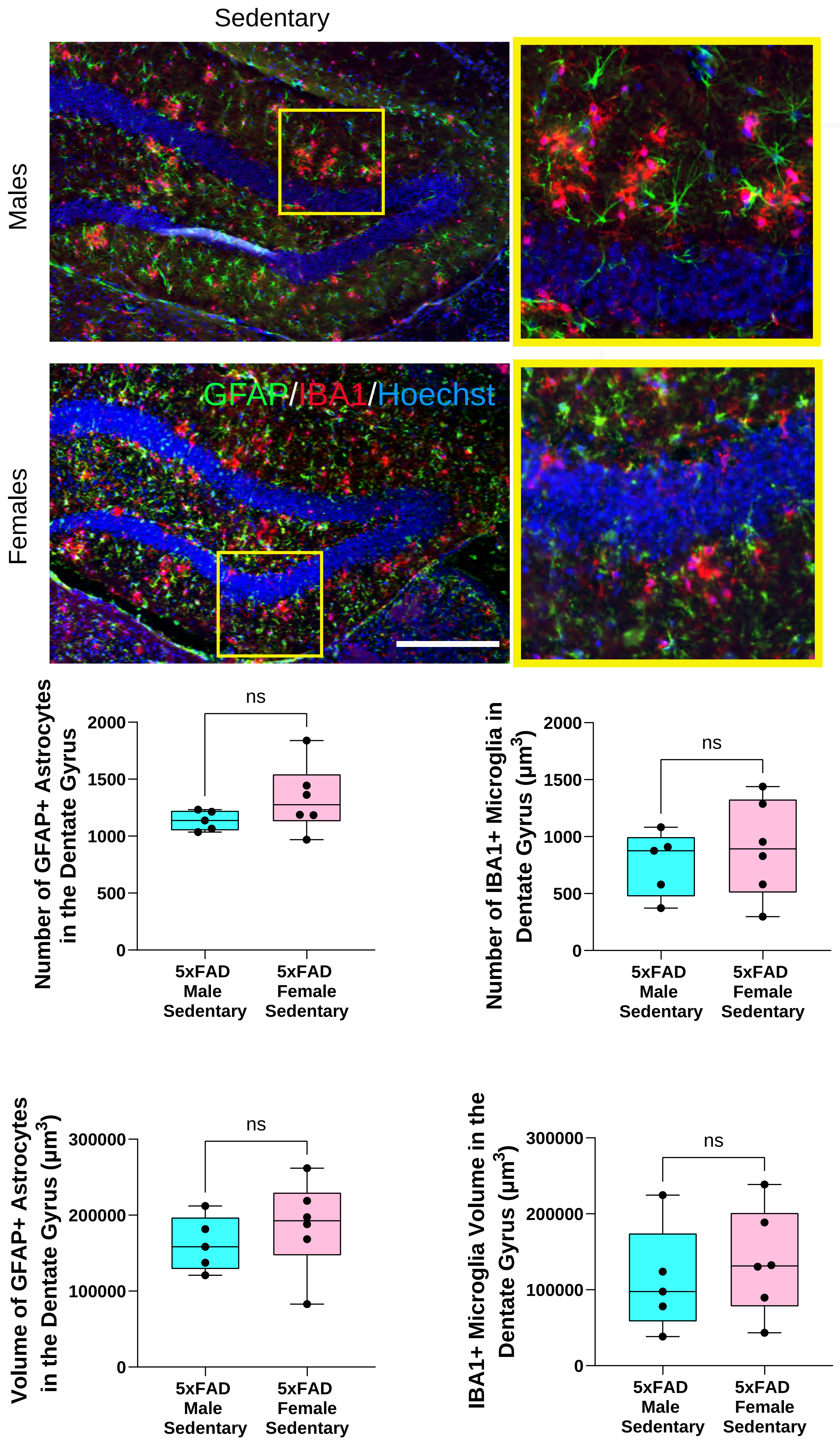

### Supplementary Figure 2 (males)

#### A. Principal component analysis of male 5xFAD hippocampal transcriptomes

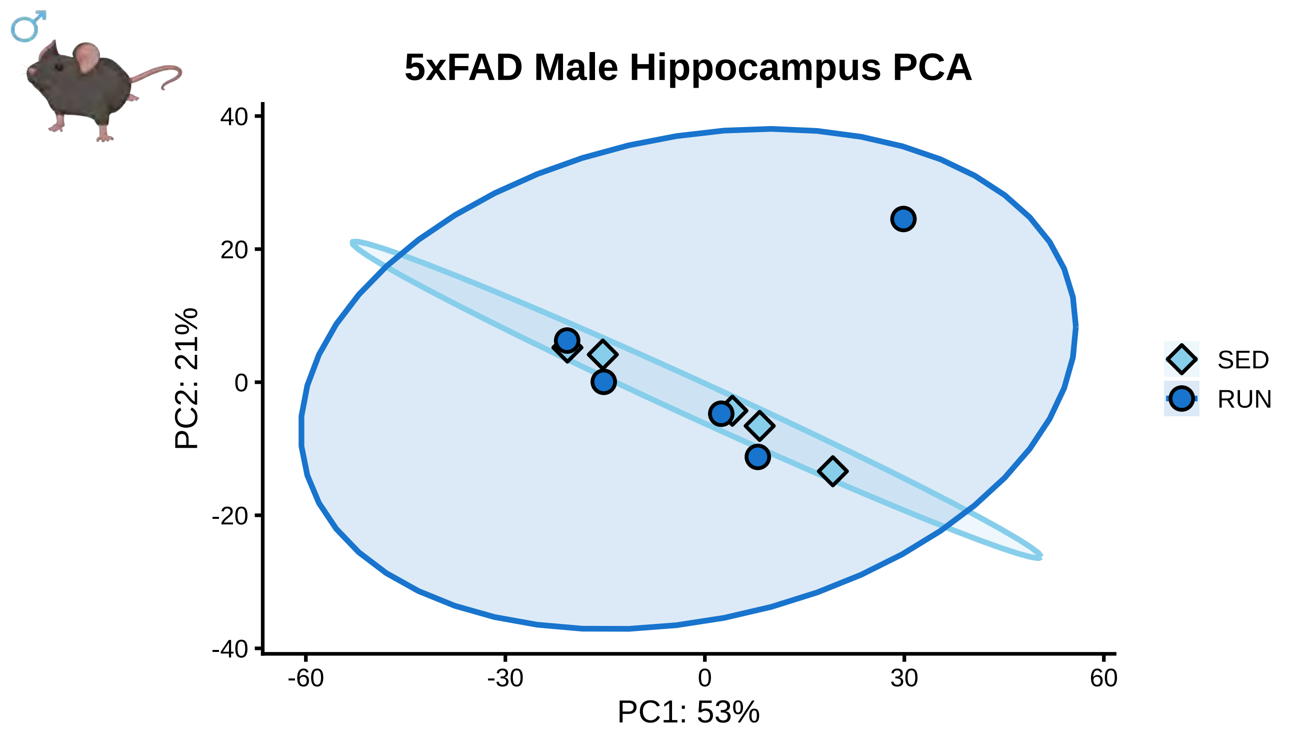


#### B. Volcano plots for differential gene expression of the male 5xFAD hippocampus

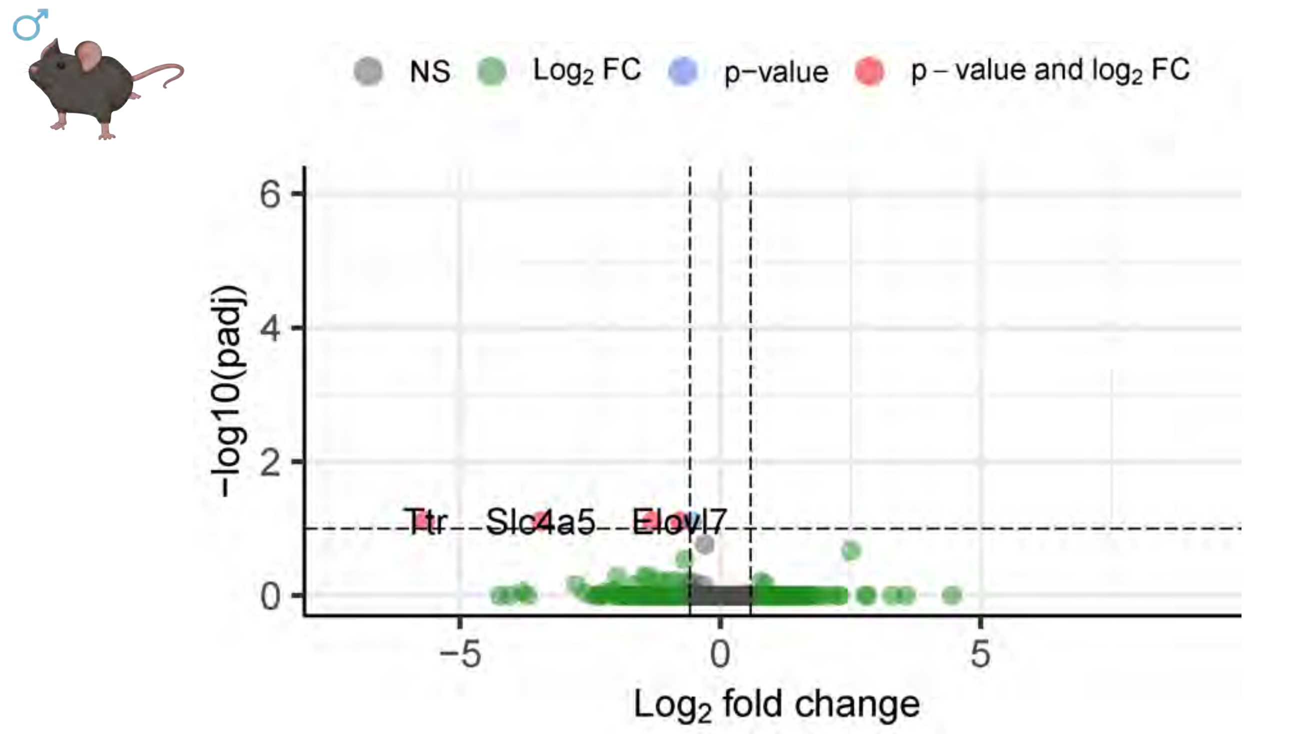


#### C. Top 30 GO Cell Compartment terms in runner male 5xFAD hippocampus

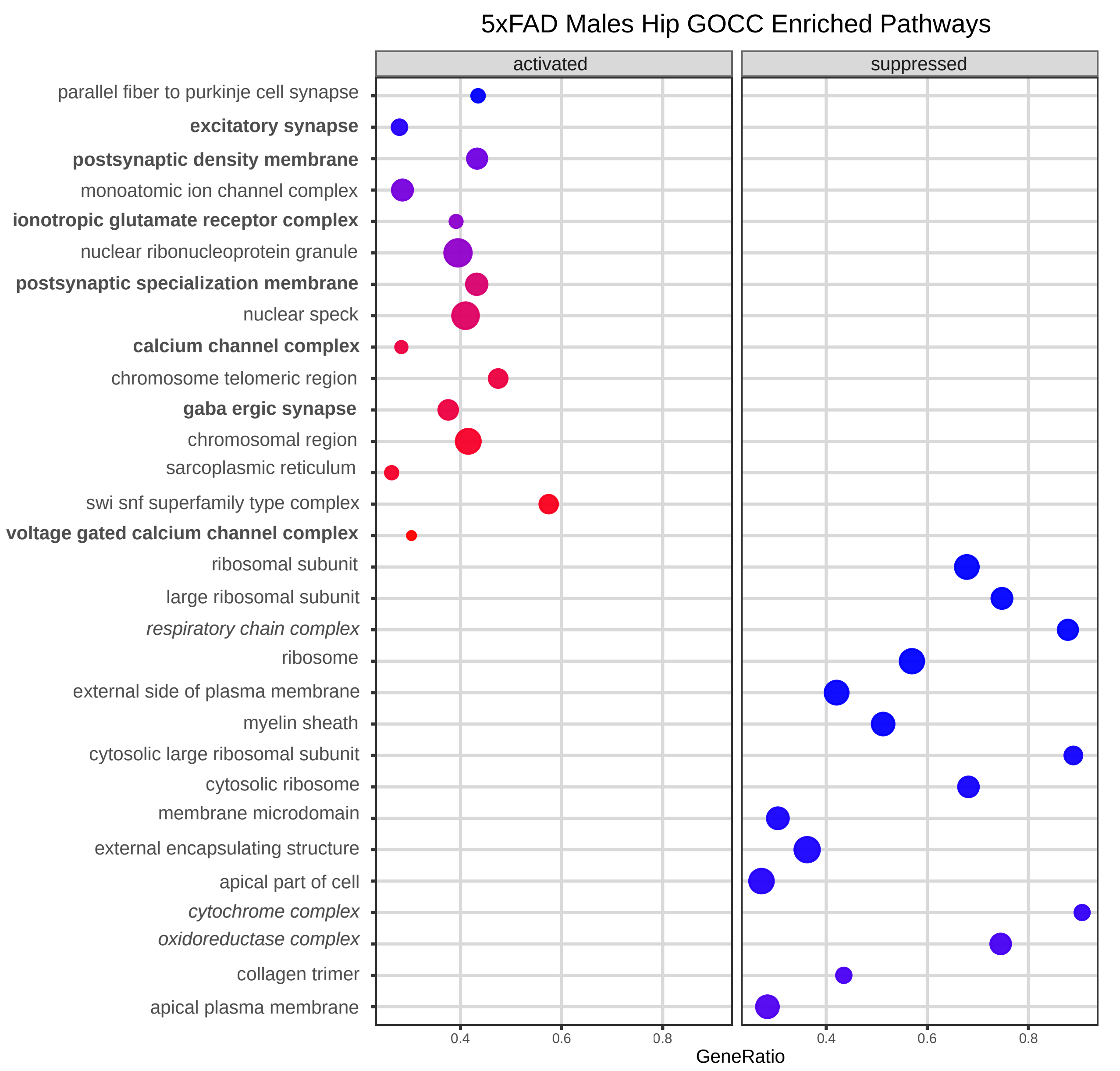


#### D. Top 30 GO Molecular Function terms in runner male 5xFAD hippocampus

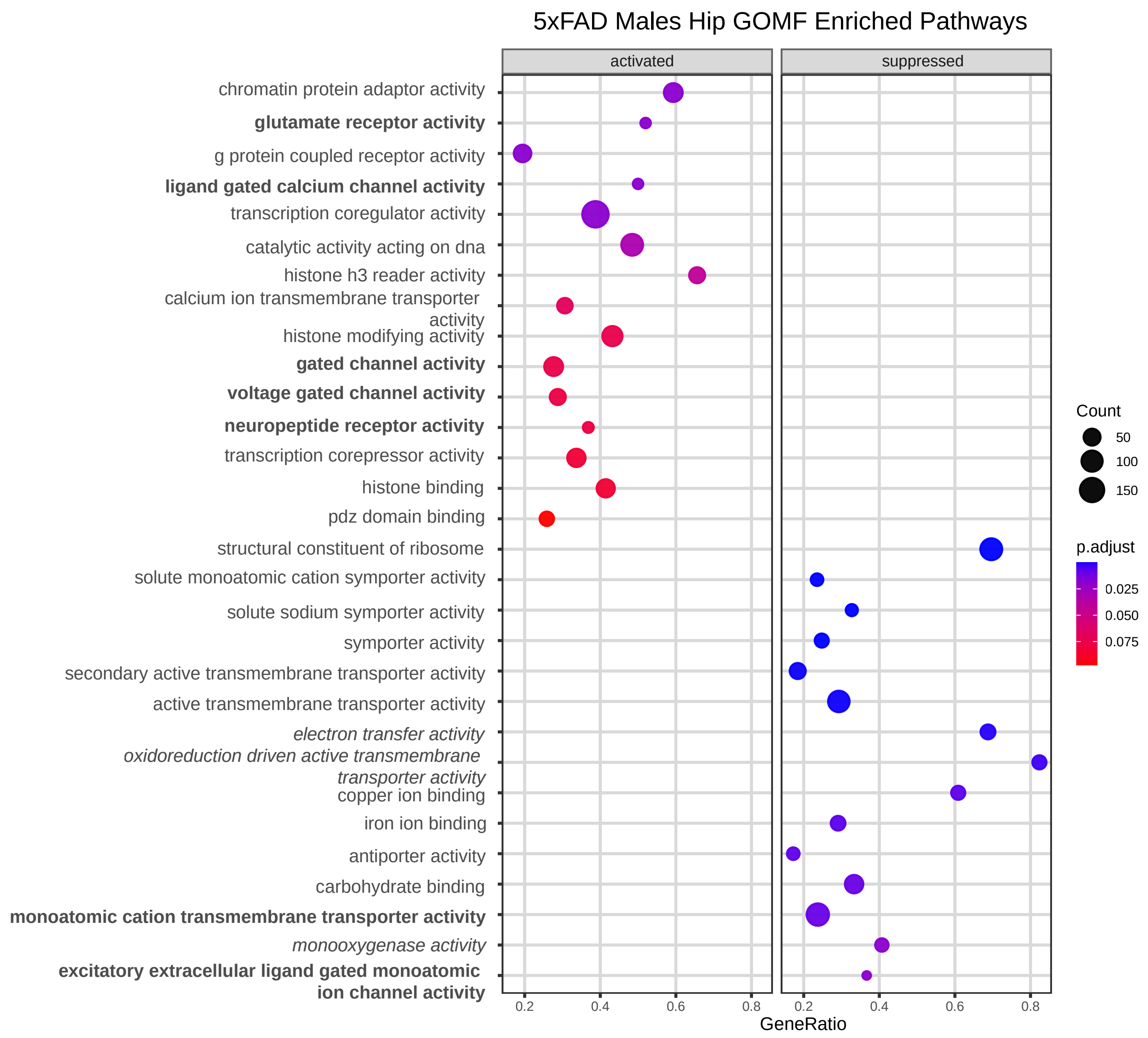


#### E. Quantification of average hippocampal astrocyte number

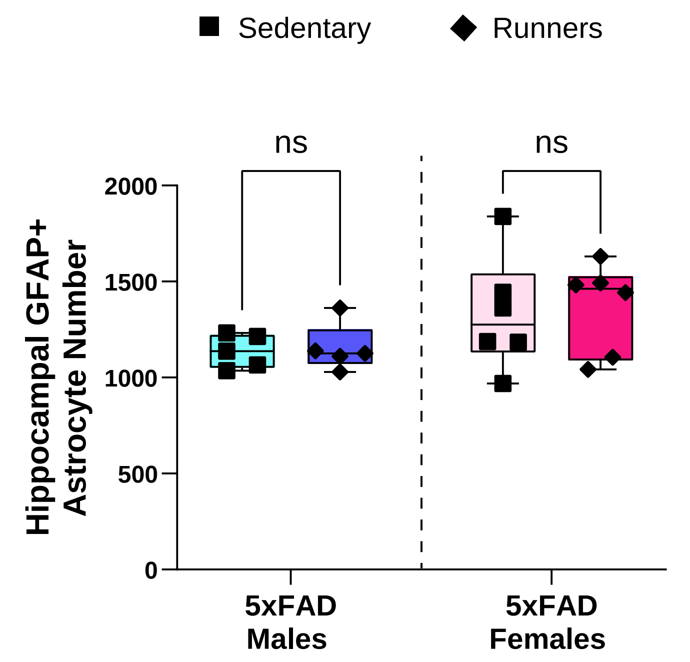


#### F. Size distribution of GFAP+ structures in the dentate gyrus of 5xFAD mice

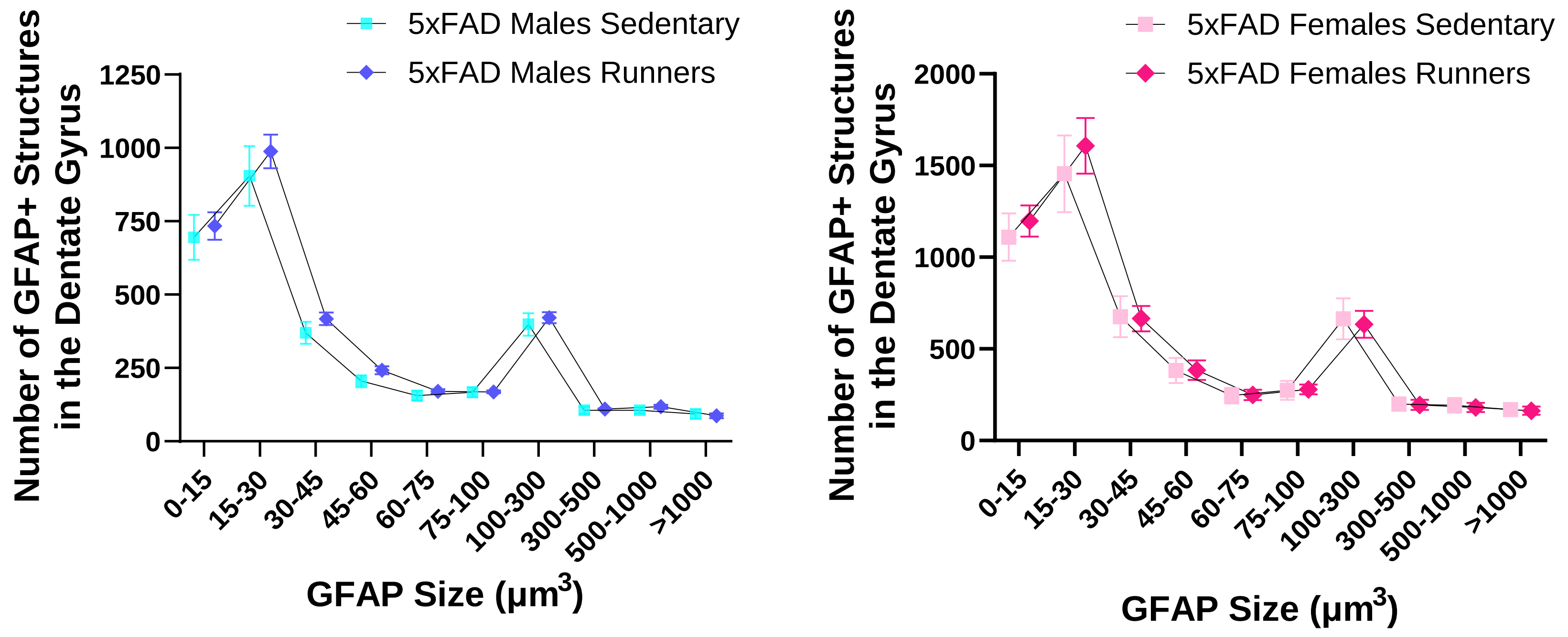


#### G. Representative images of microglial morphometric states in the 5xFAD dentate gyrus

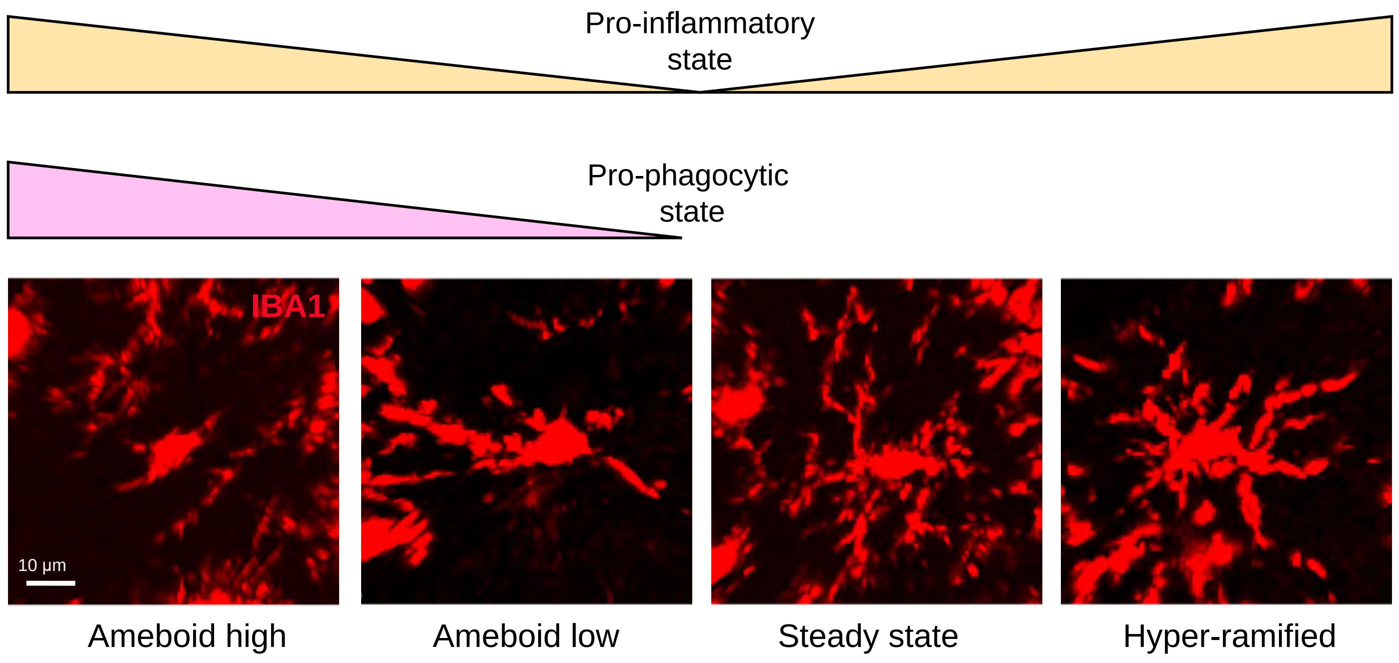


#### H. Quantification of IBA+ microglia shape

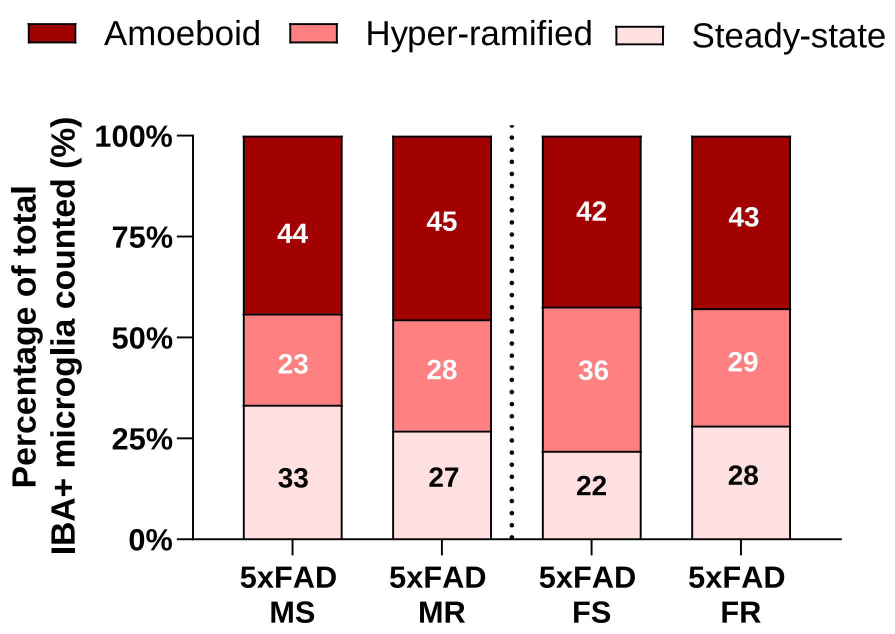

### Supplementary Figure 3 (females)

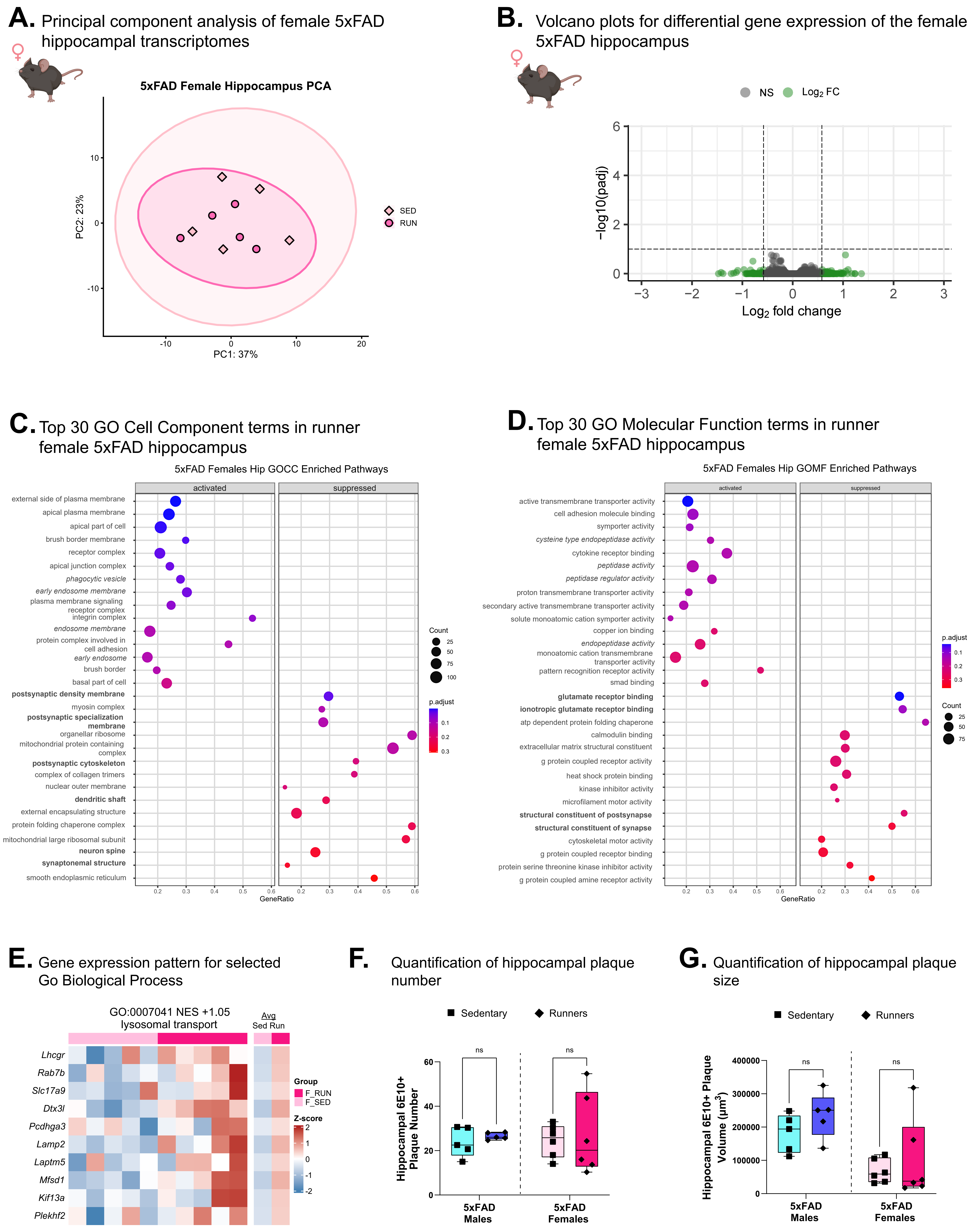

### Supplementary Figure 4 (Behavior)

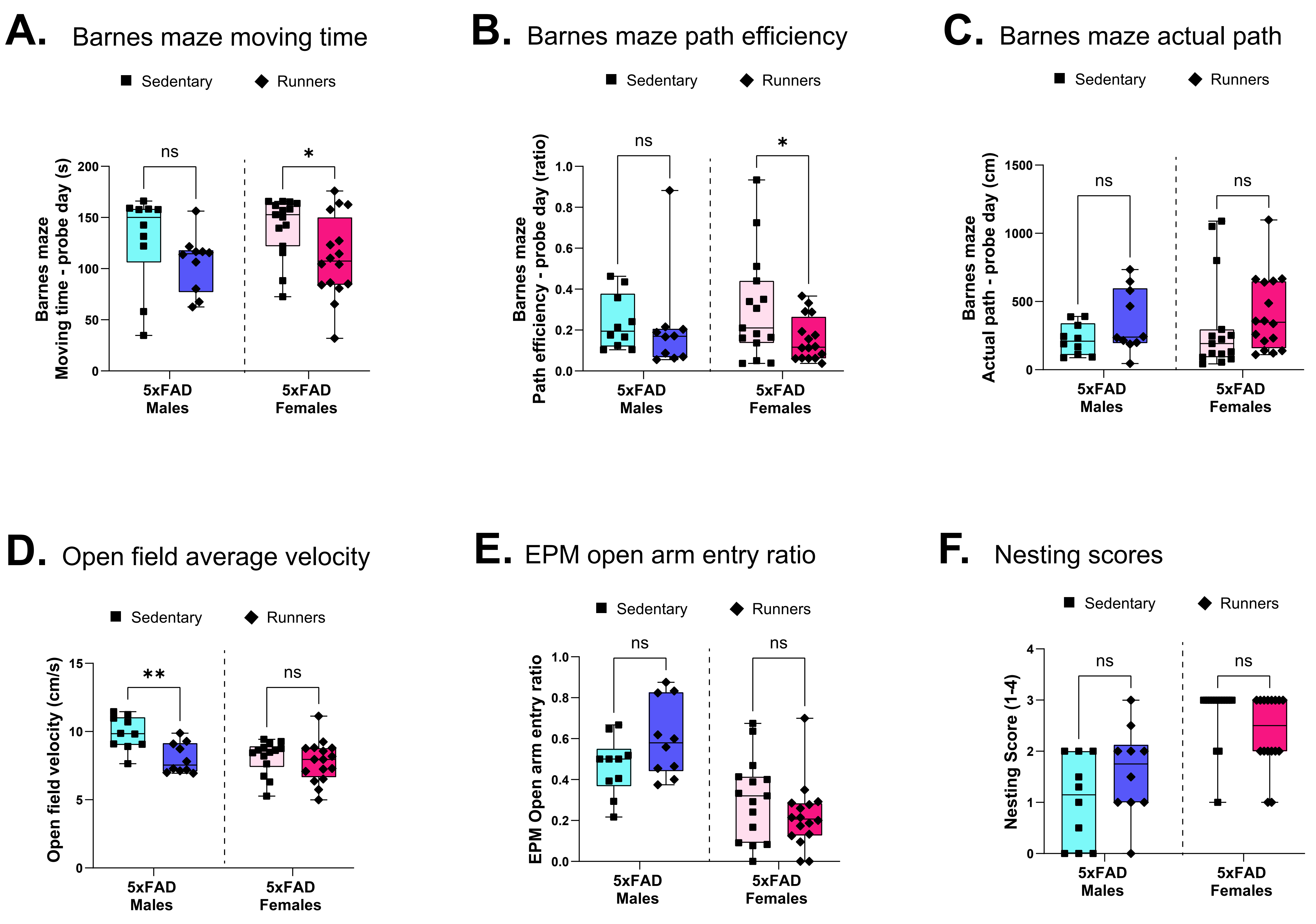

### Supplementary Figure 5 (iWAT)

**A.** iWAT PCA sex + age X exercise

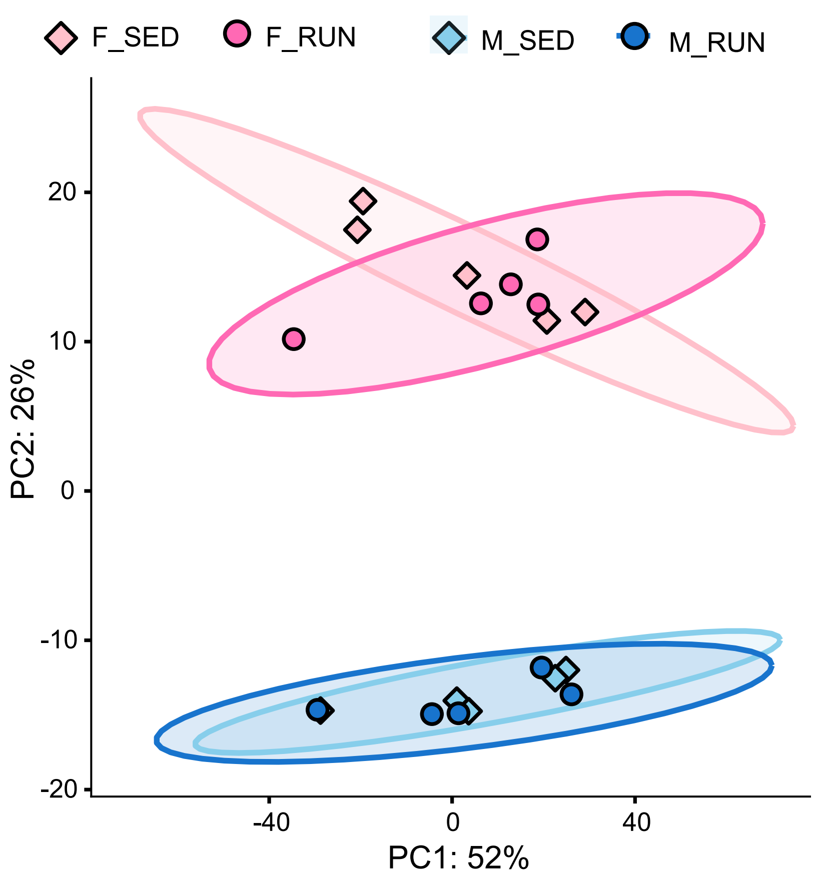


**B.** Exercise-induced transcriptional changes in iWAT by sex

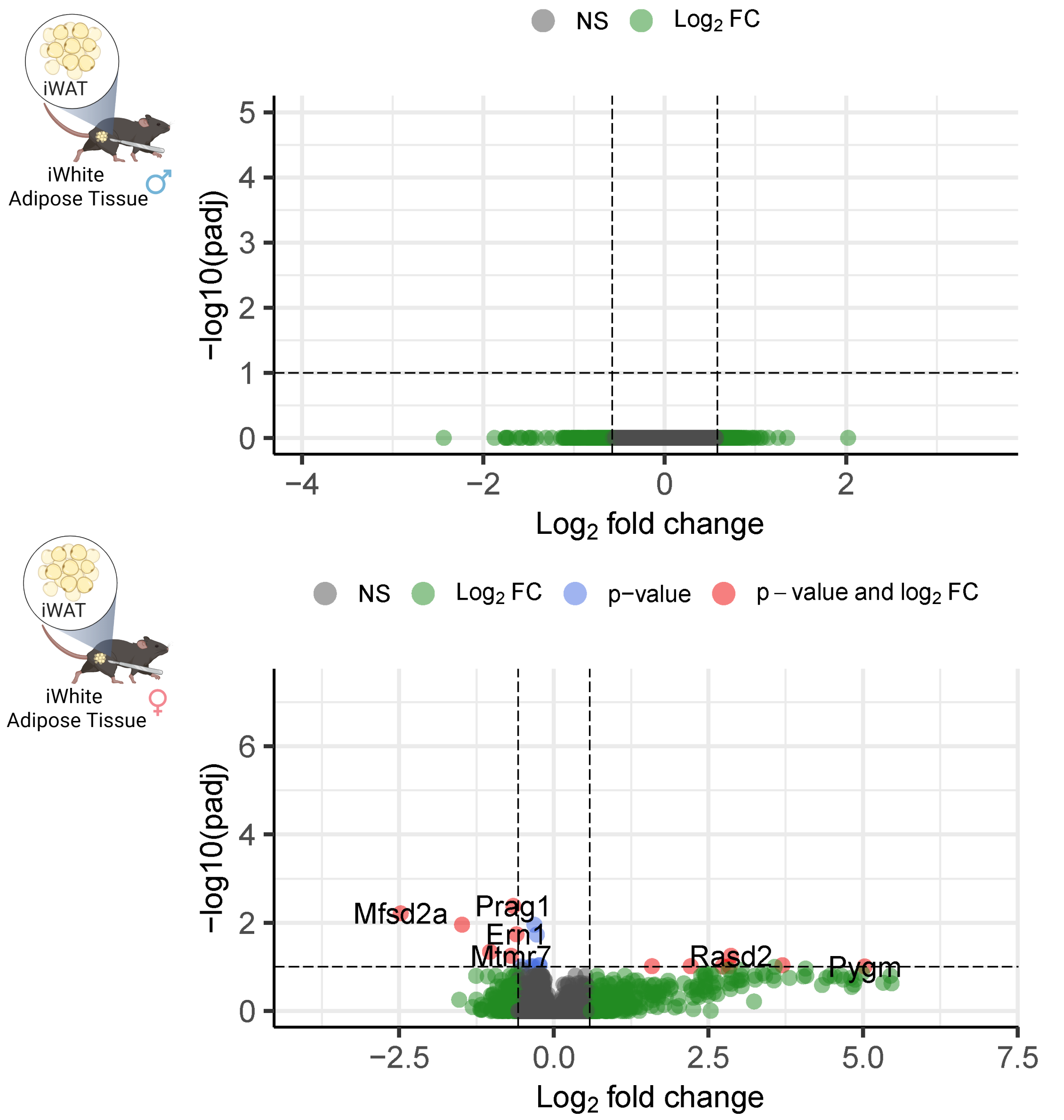


**C.** Top 20 GO biological processes enriched in male 5xFAD iWAT following voluntary wheel running

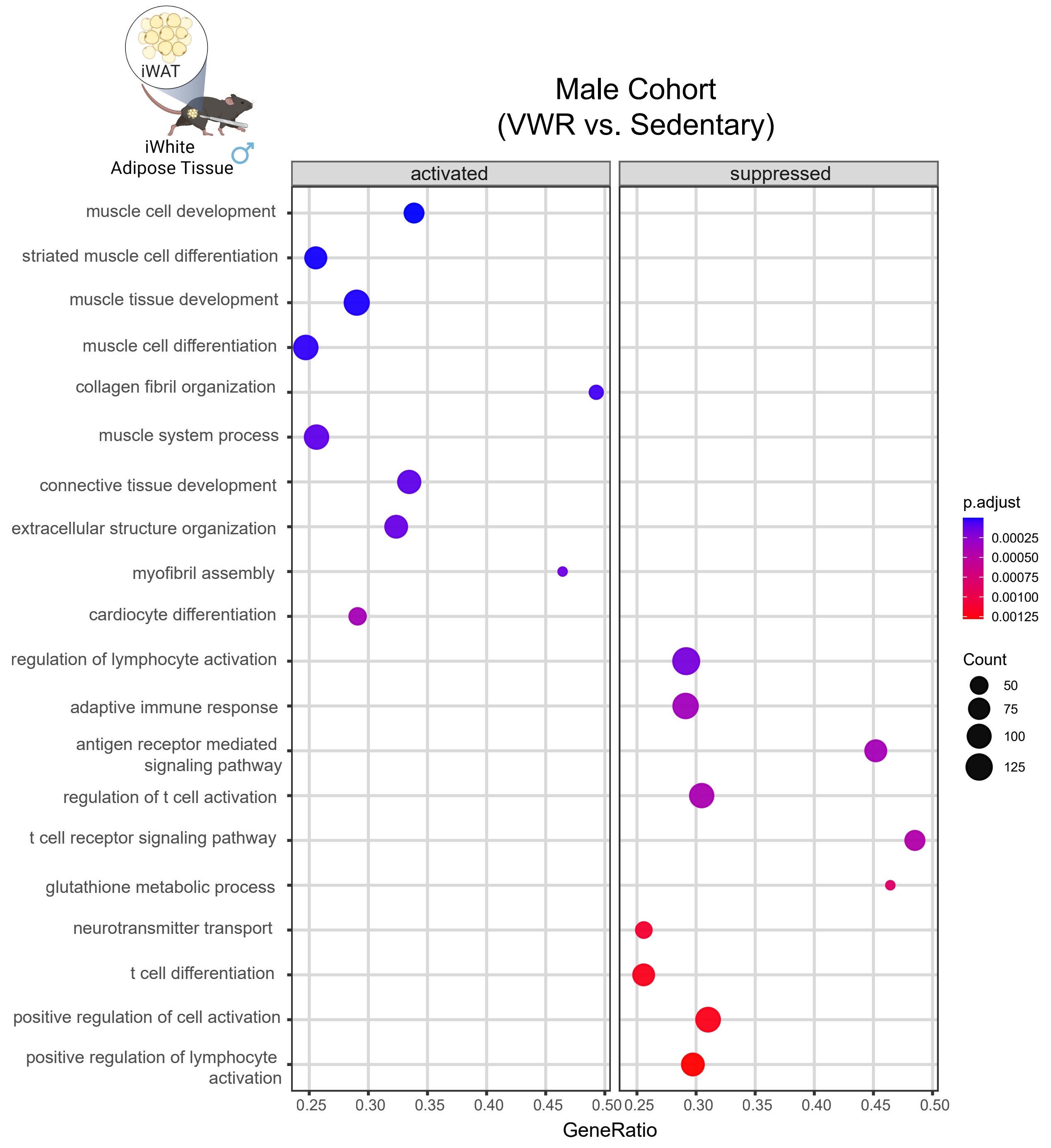


**D.** Top 20 GO biological processes enriched in female 5xFAD iWAT following voluntary wheel running

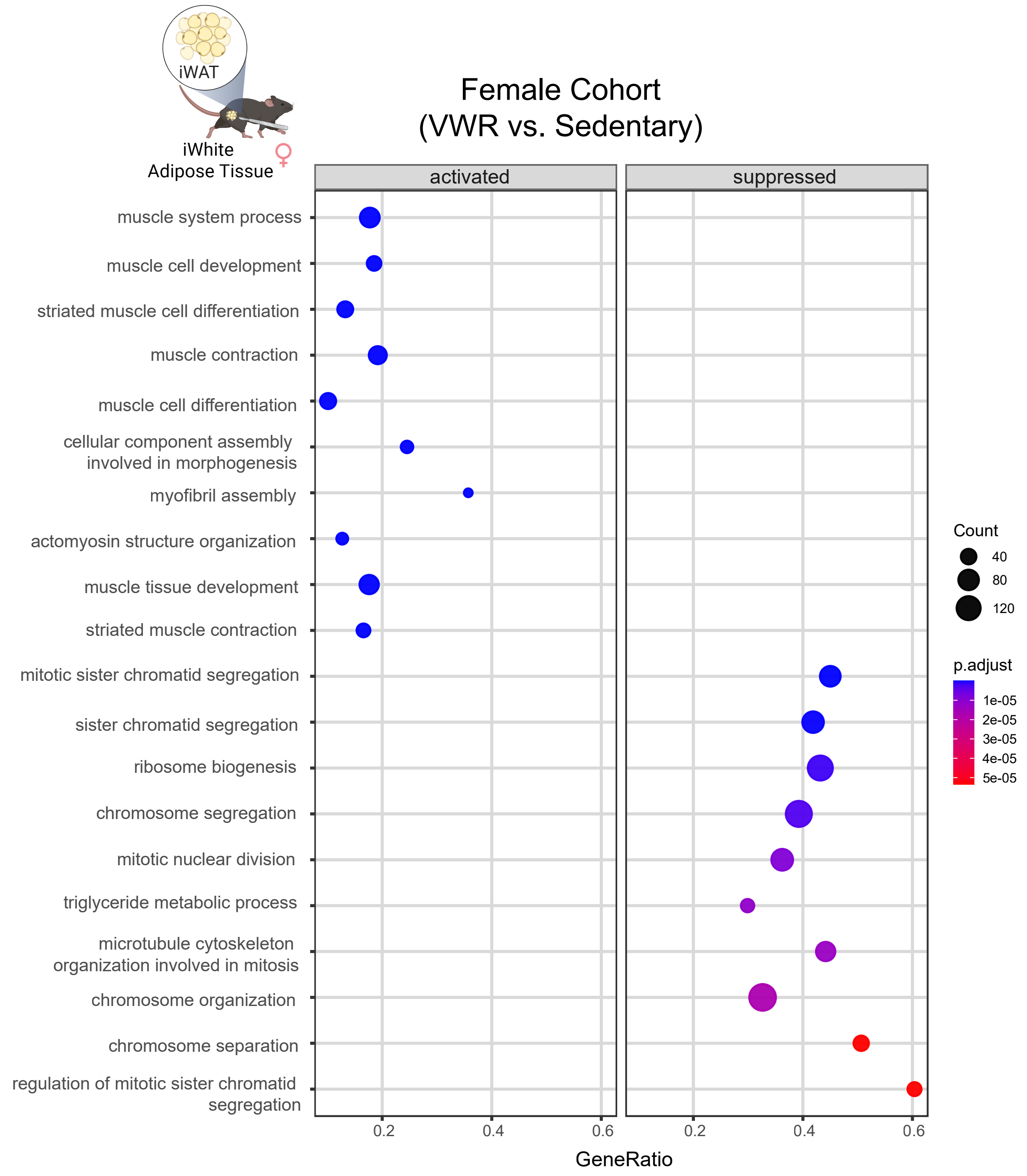
